# The kinetoplastid kinetochore protein KKT23 acetyltransferase is a structural homolog of GCN5 that acetylates the histone H2A C-terminal tail

**DOI:** 10.1101/2024.08.16.608276

**Authors:** Patryk Ludzia, Midori Ishii, Gauri Deák, Christos Spanos, Marcus D. Wilson, Christina Redfield, Bungo Akiyoshi

**Affiliations:** Department of Biochemistry, University of Oxford, Oxford, OX1 3QU, UK; Wellcome Centre for Cell Biology, University of Edinburgh, Edinburgh, EH9 3BF, UK

**Keywords:** kinetochore, kinetoplastid, *Trypanosoma brucei*, KKT23, histones, acetyltransferase, acetylation, X-ray crystallography, NMR spectroscopy

## Abstract

The kinetochore is the macromolecular protein machine that drives chromosome segregation in eukaryotes. In an evolutionarily divergent group of organisms called kinetoplastids, kinetochores are built using a unique set of proteins (KKT1–25 and KKIP1–12). KKT23 is a constitutively localised kinetochore protein containing a C-terminal acetyltransferase domain of unknown function. Here, using X-ray crystallography and NMR spectroscopy, we have determined the structure and dynamics of the KKT23 acetyltransferase domain from *Trypanosoma brucei* and found that it is structurally similar to the GCN5 histone acetyltransferase domain. We find that KKT23 can acetylate the C-terminal tail of histone H2A and that knockdown of KKT23 results in decreased H2A acetylation levels in *T. brucei*. Finally, we have determined the crystal structure of the N-terminal region of KKT23 and shown that it interacts with KKT22. Our study provides important insights into the structure and function of the unique kinetochore acetyltransferase in trypanosomes.

## Introduction

During cell division, genetic material must be replicated and segregated through the process of chromosome segregation [1]. This process is facilitated by the kinetochore complex, whose components are widely conserved across eukaryotes [2, 3]. However, a unicellular group of eukaryotes called kinetoplastids has a unique set of kinetochore proteins that lack a significant similarity to conventional kinetochore proteins. These proteins were discovered in *Trypanosoma brucei* and consist of kinetoplastid kinetochore proteins (KKT1–25) and KKT-interacting proteins (KKIP1–12) [4–8]. In this study, we focus on KKT23, which was identified as a protein that co-purified with a constitutively localised kinetochore protein KKT3 [6]. Fluorescence microscopy of YFP-KKT23 revealed that it too localises at the kinetochore throughout the cell cycle, and its immunoprecipitation from trypanosomes revealed co-purification of many kinetochore proteins, including KKT3 and KKT22 [6]. An RNAi-mediated knockdown of KKT23 resulted in growth defect, showing that KKT23 is a vital component of the kinetochore complex in kinetoplastids [9]. Intriguingly, KKT23 has a predicted GCN5-related N-acetyltransferase (GNAT) domain in its C-terminal region (Figure 1A) that is highly conserved among kinetoplastids [6]. Although there are acetyltransferases that regulate kinetochore functions in humans (e.g. MYST family acetyltransferases called TIP60 and KAT7), they are not considered as structural components of the kinetochore complex [10, 11]. Therefore, the presence of a GNAT protein is a unique feature of kinetochores in kinetoplastids. The function or structure of KKT23 remain unknown.

**Figure 1.**
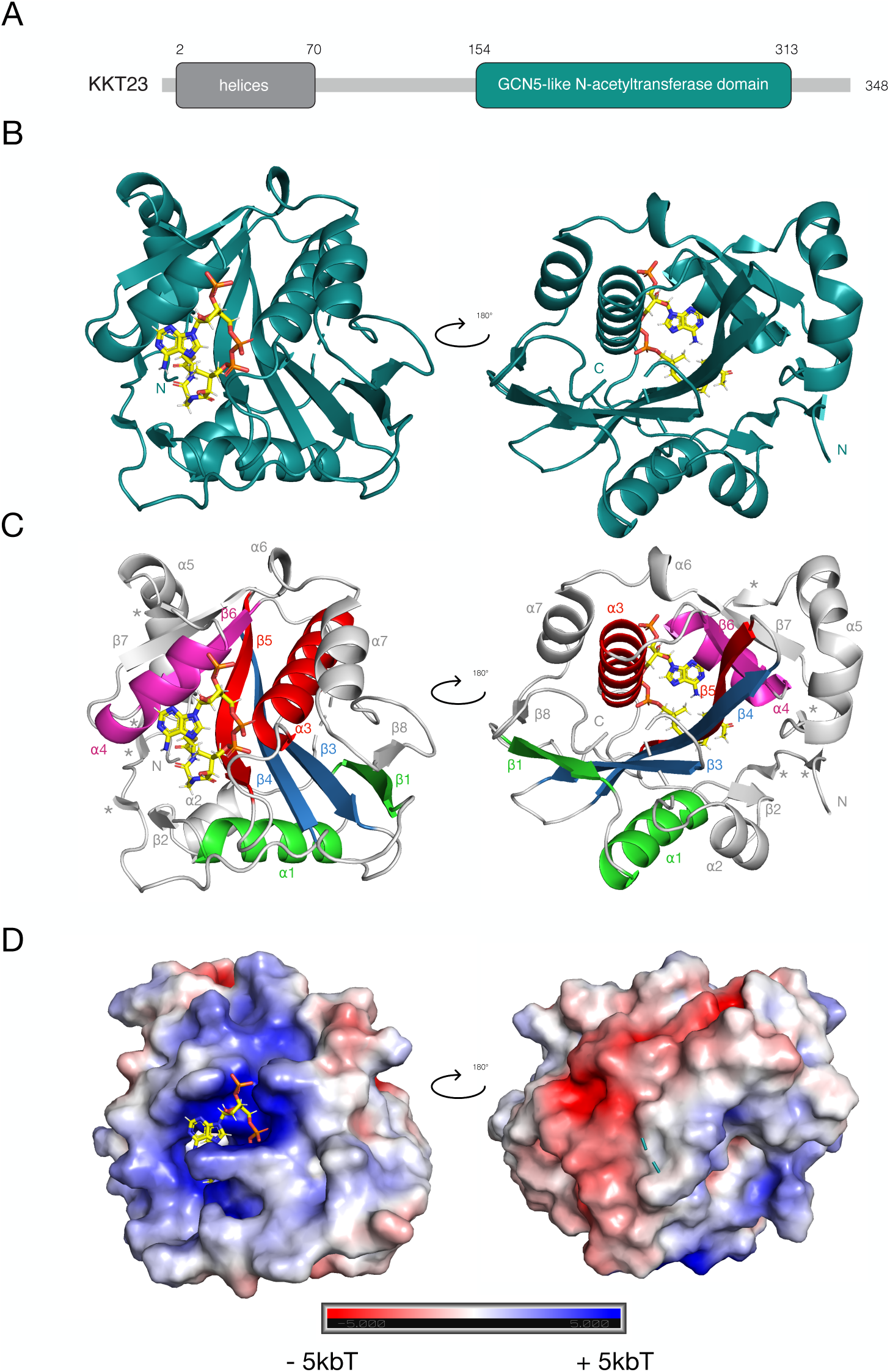
Crystal structure of KKT23^125–348^ bound to acetyl coenzyme A. A. Schematic representation of the KKT23 sequence. The N-terminal region of KKT23 is predicted to be mostly helical, while the C-terminal region is predicted to have a GCN5-related N-acetyltransferase domain fold [6]. See also Figure S1. B. Ribbon model of the *Trypanosoma brucei* KKT23^125–348^ crystal structure. Acetyl coenzyme A, shown as sticks is coloured by the elements (C - yellow, N - blue, O - red, P-orange). The secondary structure elements are labelled in (C). N and C represent the N- and C-termini, respectively. C. Motifs of the GNAT domain are highlighted in the crystal structure using different colours: motif A (red), B (magenta), C (green) and D (blue). The core of the enzyme is composed of motifs A and D. The four asterisks represent short β-strands, which were not included in the numbering of ⍺/β elements. See also Data S1. D. The surface electrostatic potential is coloured from red (acidic) to blue (basic), -5 kbT to +5 kbT, as calculated by APBS electrostatic plugin in Pymol [40]. See also Data S1.

In this report, we present structural and functional analysis of KKT23 in *Trypanosoma brucei*. We have revealed that the structure of its acetyltransferase domain is most similar to the GCN5 histone acetyltransferase. We also show that KKT23 can acetylate nucleosomes *in vitro* and have identified specific sites on histone H2A that are acetylated by KKT23. Notably, we have found that knockdown of KKT23 results in reduced acetylation levels on histone H2A in *T. brucei*. Our study has also revealed that KKT23 forms a complex with other kinetochore proteins, including KKT3 and KKT22, via its N-terminal region. These results provide valuable insights into the structure and function of the KKT23 acetyltransferase located at kinetoplastid kinetochores.

## Results

### C-terminal region of KKT23 has an acetyltransferase domain

Our sequence analysis of KKT23 predicted a helical region near the N-terminus (4–66), followed by a stretch of unstructured residues (67–128) (Figure S1). The C-terminal region of KKT23 (146– 313) is predicted to be a mixture of ⍺-helices and β-strands, consistent with the GNAT domain prediction in that region (Figure 1A, Figure S1A). We first employed Size Exclusion Chromatography with Multi-Angle Light Scattering (SEC-MALS) and Analytical Ultracentrifugation (AUC) and found that the recombinant full-length protein is monomeric (Figure S2). To determine a high-resolution structure of KKT23, we next used X-ray crystallography. However, no crystal for the full-length protein was obtained. We therefore aimed to solve the structure of the acetyltransferase domain (KKT23^125–348^) in the presence of a known cofactor of acetyltransferases, acetyl coenzyme A (acetyl-CoA). These efforts led to robust crystal formation, with the best crystals diffracting up to 1.8 Å resolution. Our attempts to solve the native structure using molecular replacement were unsuccessful. Instead, the X-ray structure of native KKT23^125–348^ was solved to 1.8 Å resolution using a selenomethionine derivative of KKT23^125–348^ (Figure 1B). The data collection and refinement statistics of both structures are summarised in Table 1.

**Table 1.**
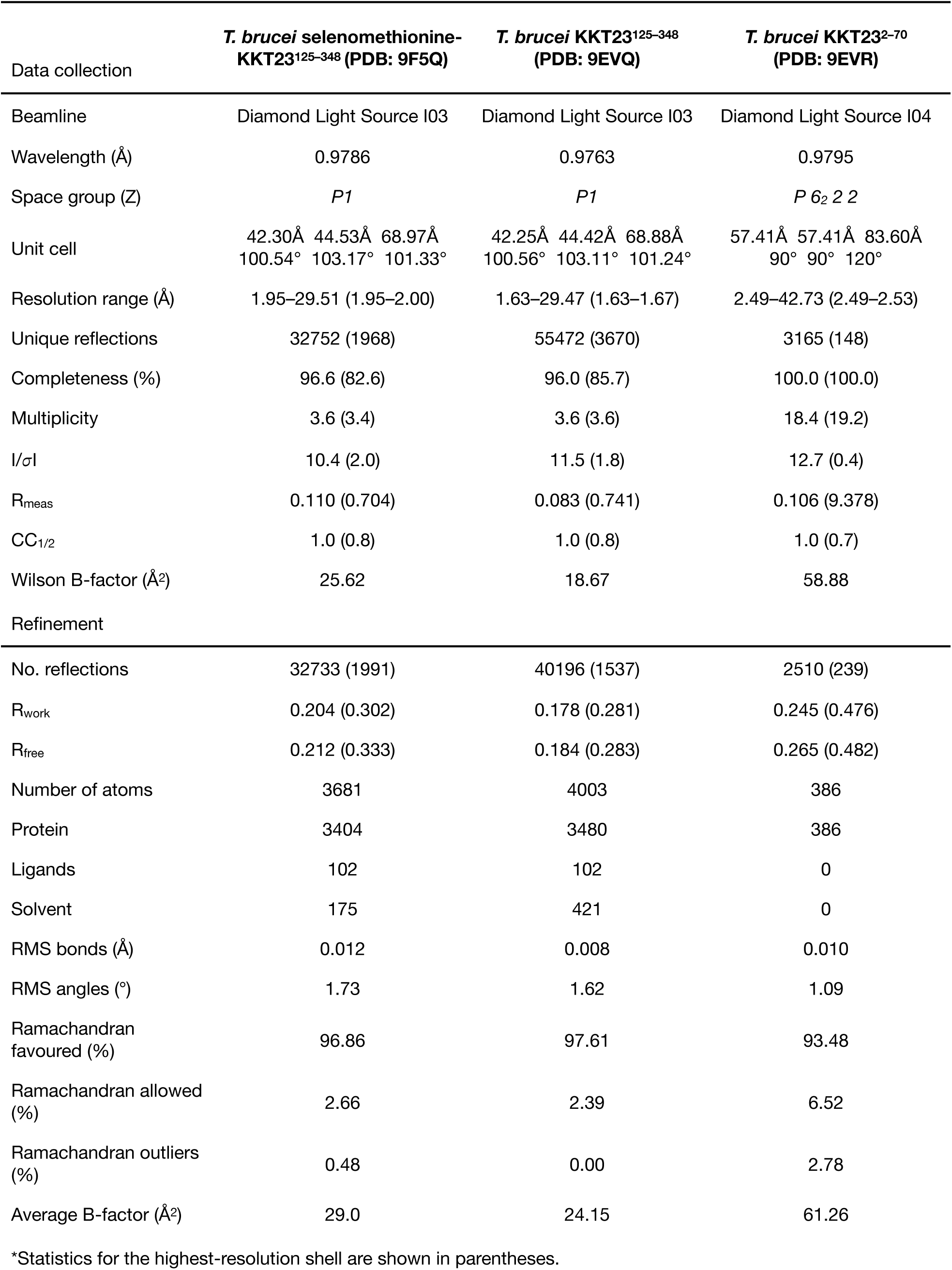
Data collection, refinement statistics*.

In the asymmetric unit of the crystal, two molecules (A and B) of KKT23^125–348^ are both bound to acetyl-CoA. In molecule A, the electron density is missing for the N-terminal residue Q125 and residues 192–200, whereas in molecule B, the N-terminal regions 125–126, 137–139 and 195–198 are missing, most likely because they are disordered and/or flexible. The overall structure of KKT23^125–348^ has a mixed ⍺/β topology containing 7 ⍺-helices and 8 β-strands organised in 4 motifs (A, B, C and D), as shown in Figure 1C and Data S1; this motif organisation is consistent with previously reported structures of N-acetyltransferases [12]. The core of the enzyme is built by a three-stranded antiparallel β-sheet (strands 3,4 and 5) that sits below ⍺-helix 3 (motifs A and D). The β-strands β3 and β4 (motif D) near the N-terminus are hydrogen-bonded to each other, most likely stabilising the core region. Another part of the protein core is a β-loop-⍺ motif built by β6 and ⍺4 (motif B). In contrast to the secondary structure prediction (Figure S1A), no helices are observed in the crystal structure between residues 125 and 150 (Figure 1B). Using DSSP Stride [13], additional short β-strands were found in the structure. Two of these strands, involving residues 129–130 and 133–134, are located in the N-terminal region. The strand between residues 129 and 130 makes contacts with region 308–309, while region 133–134 is hydrogen bonded to β2 as shown in Figure 1C and Figure S5. The hydrogen bonds formed by these short strands stabilise the mostly unstructured N-terminal region, which explains its well defined electron density. Finally, the overall electrostatic potential of the protein surface shows a mixture of basic and acidic regions with the most positively charged region binding to acetyl-CoA (Figure 1D, Data S1).

### Acetyl-CoA is coordinated by residues located in motif A of the KKT23 GNAT domain

Based on the analysis using PDBePISA [14] and the crystal structure, acetyl-CoA is coordinated by neighbouring residues through both hydrogen bonds and hydrophobic interactions (Figure 2, Figure S4). Residues that were found to form hydrogen bonds with acetyl-CoA in both molecules of the asymmetric unit include F239, T241, K247, G249, F250, G251, R252 and H284. In molecules B, additional hydrogen bonds were observed with H248 and R333. F239 and T241 are located on β5; K247 occupies the loop region preceding ⍺3, G249, F250, G251 and R252 are located on ⍺3, while H284 is located on ⍺4. Most of the residues that form hydrogen bonds with acetyl-CoA are conserved among kinetoplastids (Figure S3) and are located within motif A, which is consistent with the previous studies of acetyltransferases within the GNAT superfamily [12, 15].

**Figure 2.**
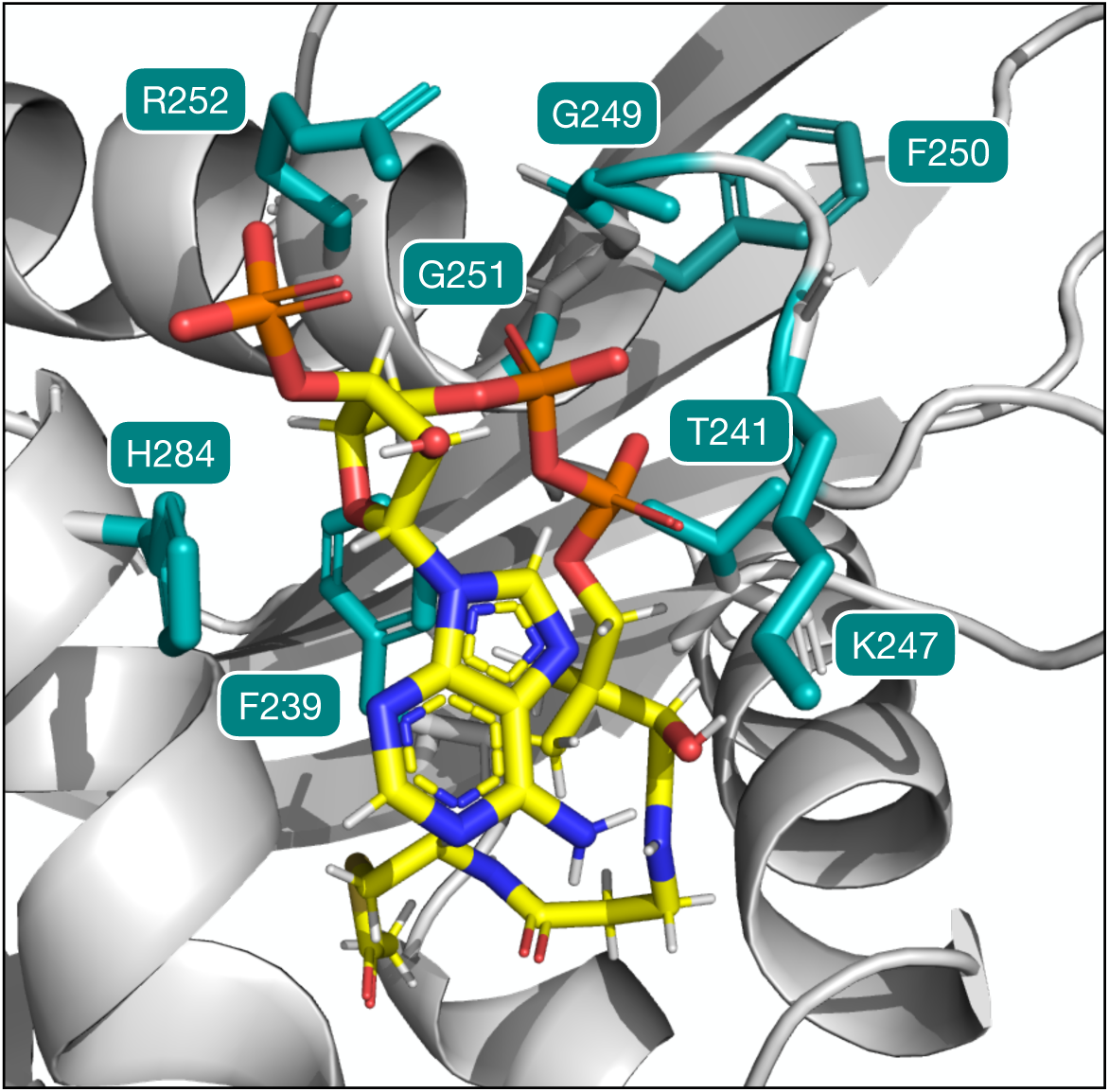
Coordination of acetyl-CoA in the KKT23^125–348^ crystal structure. Residues within the KKT23^125–348^ crystal structure (teal) that are coordinating acetyl coenzyme A using hydrogen bonds are shown. Acetyl coenzyme A is coloured by the elements. See also Figure S4.

### Solution structure of KKT23^125–348^ resembles the crystal structure

To characterise the structure and dynamics of KKT23^125–348^ in solution, we employed NMR spectroscopy. Using isotope-labelled KKT23^125–348^, we have obtained a detailed backbone resonance assignment in the presence of acetyl-CoA. Using the obtained chemical shifts, TALOS-N was used to predict the secondary structure elements in KKT23^125–348^ in solution [16], showing a mixture of ⍺-helices and β-strands (Figure 3A, Data S2). Overall, the secondary structure elements identified by TALOS-N are in excellent agreement with those identified in the crystal structure (Figure 3B, Data S2). It is noteworthy that as in the crystal structure, the NMR analysis confirmed the lack of the predicted helix between residues 125–150. The β-strands involving residues 129– 130 and 308–309 have not been predicted by TALOS-N. However, this can be explained by the lack of assignments for residues in both regions. It is likely that peaks corresponding to these residues are broadened and this may indicate conformational exchange on a slower timescale for this β-sheet in solution. Additionally, TALOS-N does predict the short β-strand involving residues 133–134 but shows no evidence for strand β2 (residues 175–177) (Figure 3A, Figure S5, Data S2).

**Figure 3.**
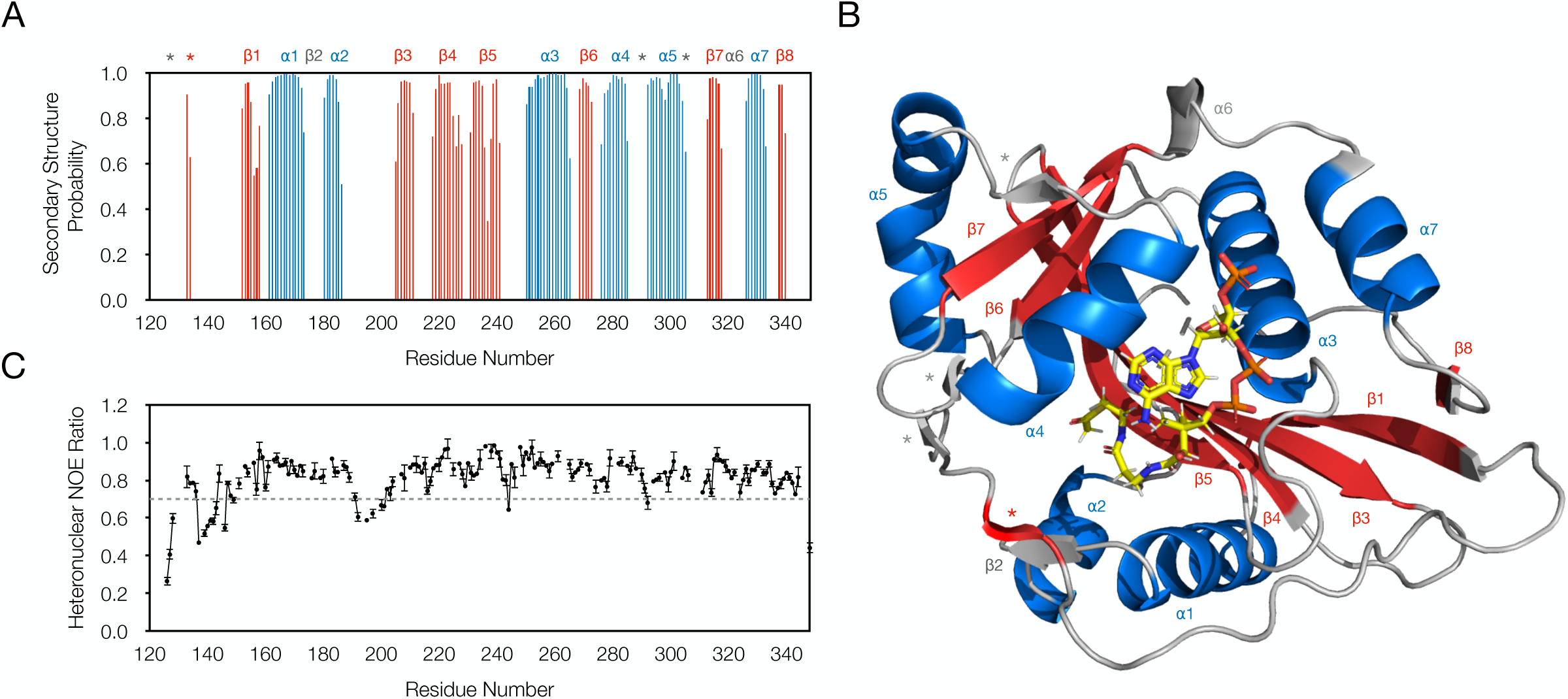
NMR analysis of KKT23^125–348^ structure and dynamics. A. The TALOS-N secondary structure analysis of KKT23^125–348^ shows a mixed ⍺/β topology. The secondary structure elements for which the probability was lower than 0.5 are not shown in the figure. See also Data S2. B. The secondary structure elements identified using TALOS-N (A) are highlighted in the crystal structure of KKT23^125–348^ showing a good agreement between the two methods. The β-strands and ⍺-helices are coloured in red and blue, respectively. See also Data S2. C. The {^1^H}-^15^N heteronuclear NOE ratios were measured and plotted against the sequence of KKT23^125–348^. Most of the residues display hetNOE ratios >0.7 indicating a rigid conformation of the protein backbone. The regions with higher flexibility (ratios <0.7) include 126–128, 137–143 and 192–201. The regions between 126–128 and 137–143 are coils according to the crystal structure, whereas no electron density is visible for residues 192–200 suggesting that all three regions are part of flexible loops. The {^1^H}-^15^N NOE errors were estimated from 500 Monte Carlo simulations using baseline noise as a measure of peak height error.

To study the dynamics of KKT23^125–348^, {^1^H}-^15^N heteronuclear NOE data were collected [17]. The measured {^1^H}-^15^N NOE ratios were high (>0.7) for almost all residues (Figure 3C). This means that the majority of the residues in KKT23^125–348^, even those located in most loop regions, have a rigid backbone conformation. The regions between 126–128, 137–143 and 192–201 show reduced hetNOE ratios suggesting a higher flexibility compared to the rest of the protein. This is consistent with the crystal structure and the analysis by TALOS-N, as no secondary structure was observed for these regions of the sequence. Moreover, elevated B-factors were observed for regions 126–128 and 137–143, and no electron density was observed for 192–200 (data not shown).

### KKT23^125–348^ is a structural homolog of GCN5 acetyltransferase domain

Our previous sequence analysis showed that KKT23 has a putative GNAT domain but did not allow us to determine its closest homologue [6]. Using a structural homology search with the distance-matrix alignment (DALI) server [18] using the crystal structure of KKT23^125–348^, we identified the human GCN5 (also known as KAT2A) histone acetyltransferase domain as the top hit (Table S1). Moreover, structural homology searches with Foldseek against the AlphaFold2-predicted structure database [19] produced similar results (data not shown). The structural alignment between KKT23^125–348^ and human GCN5 revealed a remarkable similarity, with an RMSD of 1.4 Å (720 C⍺ atoms) (Figure 4A, left panel). Notable differences between the two structures include the additional ⍺-helix (⍺7) and β-strand (β8) in the crystal structure of KKT23^125–348^, which are highly conserved among kinetoplastids but are not part of the consensus GNAT domain fold (Figure S3). To highlight structural differences with other classes of the GNAT family, the crystal structure of KKT23^125–348^ was aligned with the next top hit found by the DALI search, *D. melanogaster* NAA80 N-terminal acetyltransferase (Figure 4A, right panel). The NAA80 is a member of the GNAT family that catalyses the transfer of an acetyl group from acetyl-CoA to the N-termini of protein substrates [20, 21]. The most significant differences between KKT23 GNAT and NAA80 are observed within motif D, which builds the core of the GNAT domain (strands β3 and β4) (Figure 4B). Taken together, our structural characterisation reveals that KKT23 has an acetyltransferase fold with the closest structural similarity to GCN5 histone acetyltransferases.

**Figure 4.**
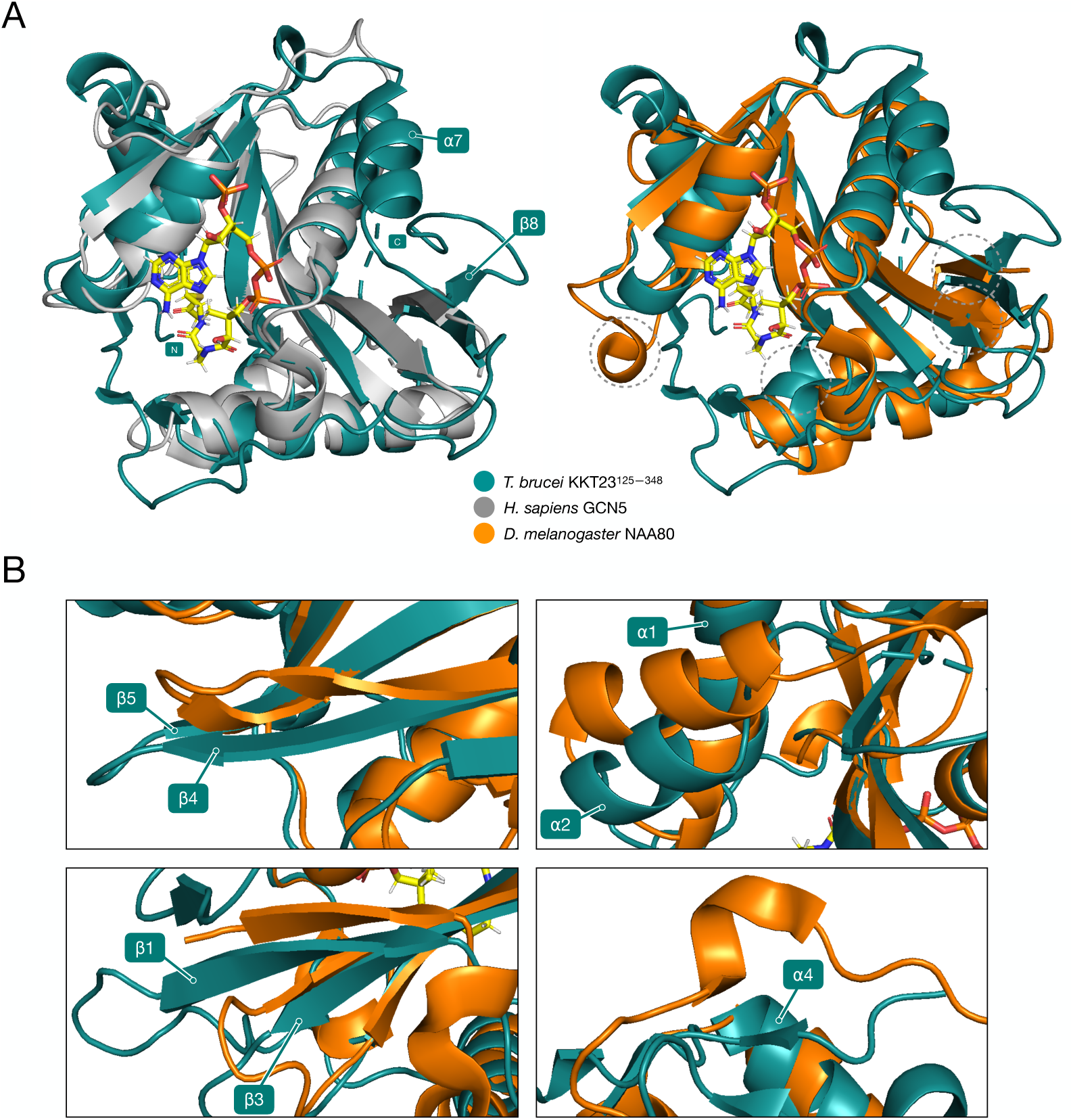
KKT23 C-terminal region is a structural homologue of GCN5. A. Superposition of the KKT23^125–348^ (teal, PDB: 9EVQ) with *Hs*GCN5 histone acetyltransferase domain (grey, PDB: 5TRM) and *Dm*NAA80 N-terminal acetyltransferase (orange, PDB: 5WJD). The structures were overlaid using *super* function in Pymol [13] with an RMSD of 1.4 Å (720 C⍺ atoms) and 1.3 Å (525 C⍺ atoms), respectively. The lower RMSD in the overlay between KKT23^125–348^ and *Dm*NAA80 can be explained by the smaller number of atoms used for the calculation. The ⍺7 and β8 in KKT23^125–348^ are not part of a canonical GNAT structure. The N and C represent the N- and C-termini. Regions of the structure highlighted with dotted circles represent close-up views shown in (B). B. The images highlight the differences between KKT23 GNAT and NAA80 N-terminal acetyltransferase domains. Most significant differences between KKT23 GNAT and NAA80 are observed within motif D, which builds the core of the GNAT domain (strands β3 and β4).

### KKT23 can acetylate *T. brucei* nucleosomes *in vitro*

GCN5 is found in numerous complexes in other well studied eukaryotes and is known to acetylate histone N-terminal tails [22–25]. The structural similarity between the KKT23 GNAT domain and the GCN5 histone acetyltransferase domain raised the possibility that KKT23 may acetylate trypanosome histones. To test this hypothesis, we performed an *in vitro* histone acetyltransferase assay. Briefly, recombinant KKT23 was incubated with acetyl-CoA and nucleosomes reconstituted using either *T. brucei* or *H. sapiens* histones [26]. Then, the reactions were analysed by immunoblots using anti-acetyl-lysine antibody. The acetylation signal was detected around the position of *T. brucei* H2A and H3 histones in the gel, but not when *H. sapiens* nucleosomes were used, suggesting that KKT23 is specific towards *T. brucei* nucleosomes (Figure 5A). To reveal the identity of the acetylated histone(s) and map the acetylated sites, mass spectrometry was employed. Gel bands corresponding to individual histones (H2A/H3, H2B, H4) were excised, digested with trypsin, and analysed by mass spectrometry. We found that lysine residues on the C-terminal tail of H2A were robustly acetylated (K119, K120, K122, K125 and K128), while acetylation on H2B, H3 and H4 was less significant (Table S2, Figure 5B). KKT23 therefore has different substrate specificity compared to GCN5 acetyltransferases in other eukaryotes that are known to preferentially acetylate histones H3 and H4 but not H2A [22–25].

**Figure 5.**
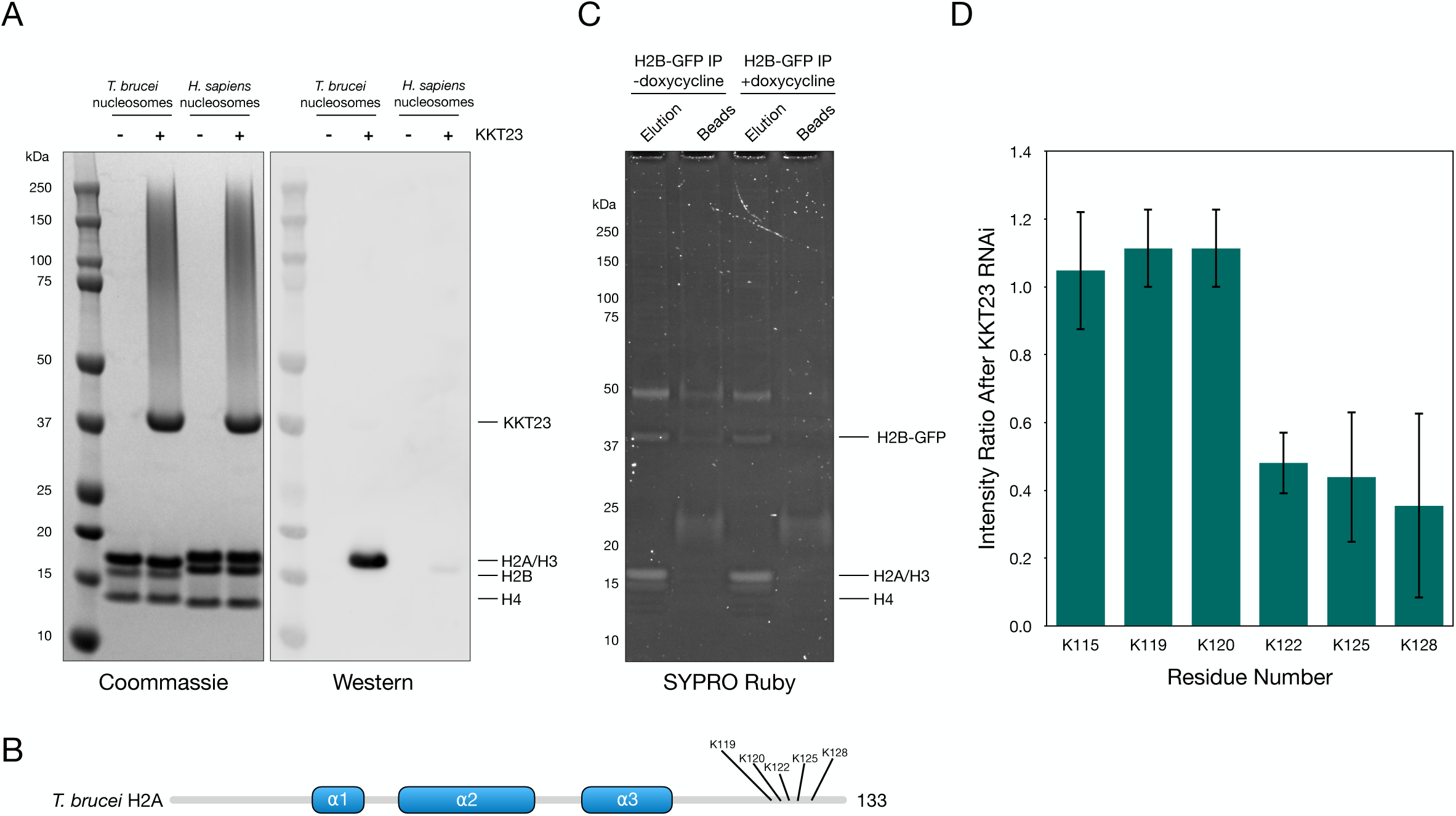
KKT23 can acetylate histone H2A. A. The SDS-PAGE gel (left panel) showing histone acetyltransferase assay on reconstituted *T. brucei* and *H. sapiens* nucleosomes using KKT23 acetyltransferase. Histones H2A and H3 from *T. brucei* and *H. sapiens* H2A and H2B could not be resolved because of their similarity in molecular weight. Panel on the right shows Western blotting analysis using the pan-acetyl-lysine antibody. Clear signal can be observed near the molecular weight of *T. brucei* H2A/H3, while no signal is observed with rest of the *T. brucei* histones or with *H. sapiens* nucleosomes, indicating that KKT23 is specific towards *T. brucei* nucleosomes. B. Schematic of *T. brucei* H2A sequence. The C-terminal lysines acetylated by KKT23 in (A) are annotated. See also Table S2. C. SDS-PAGE gel stained with SYPRO Ruby showing immunoprecipitation of H2B-GFP from *T. brucei* cells subjected to KKT23 RNAi (+ doxycycline). Control experiment was done using the same cell line but without inducing RNAi (- doxycycline). Histone H2B fused to GFP runs ∼37 kDa and co-purifies with core histones. Tubulin contamination can be observed at ∼50 kDa. Cell line, BAP2596. D. Mass spectrometry analysis of the histone H2A acetylated sites identified in the procyclic form of *T. brucei* cells, 24h after induction of KKT23 RNAi. Intensities of the acetylated peptides were normalised against intensities of total H2A in control and RNAi samples individually. Immunoprecipitation and mass spectrometry were performed three times. See also Table S3.

### KKT23 depletion leads to reduction of H2A C-terminal acetylation in *T. brucei*

Previous studies established that the C-terminal tail of histone H2A is hyperacetylated in trypanosomatids, a feature that is not found in human H2A [27–31], but the enzyme responsible for this acetylation was unknown [32]. To test a possibility that KKT23 contributes to the H2A hyperacetylation, we analysed histone acetylation levels by immunoprecipitating histone H2B-GFP from *T. brucei* cells. Using fluorescence microscopy, we confirmed that H2B-GFP was distributed in the nuclei (data not shown) and that other core histones co-purified with it (Figure 5C). We previously showed that the KKT23 protein level was reduced 2 days after RNAi induction, while the growth retardation was evident only after 4 days [9]. The acetylation levels of K122, K125, and K128 were reduced by around half after depleting KKT23 with RNAi for 2 days, while those of K115, K119, K120 did not change (Figure 5C, D; Table S3). These results show that the hyperacetylation of the histone H2A C-terminal tail partly depends on KKT23 *in vivo*.

### KKT23 forms a complex with KKT22 and KKT3

Our previous immunoprecipitation experiments suggest that KKT23, KKT22, and KKT3 are closely associated [6]. Furthermore, RNAi-mediated knockdown of KKT3 causes localisation defects for KKT22 and KKT23, while depletion of KKT22 or KKT23 did not affect KKT3 localisation [9]. To test whether KKT23, KKT22, and KKT3 directly interact, these proteins were co-expressed in insect cells (Sf9) and pulled down using FLAG-KKT23 (Figure 6A). Indeed, FLAG-KKT23 co-purified with KKT22 and KKT3 confirming a direct interaction between these proteins (Figure 6A). To dissect the hierarchy of these interactions, we repeated the experiment by omitting either KKT3 or KKT22 expression. KKT3 did not significantly co-purify with FLAG-KKT23 in the absence of KKT22 expression (Figure 6B), while KKT22 co-purified with FLAG-KKT23 even when KKT3 was absent (Figure 6C). These results suggest that KKT23 directly interacts with KKT22 and that KKT22 is required to reconstitute the complex composed of KKT3, KKT22 and KKT23 in this assay. Consistent with this possibility, KKT3 co-purified with FLAG-KKT22, but not with FLAG-KKT23 (Figure 6E). We next conducted crosslinking experiments on the purified KKT3-KKT22-KKT23 complex using BS^3^ and analysed the samples by mass spectrometry (Figure 6F). The results revealed multiple crosslinks between KKT22 and KKT23, but none were detected between KKT3 and KKT22 or KKT23. It is worth noting that numerous crosslinks were detected between the KKT23 N-terminal region and the acetyltransferase domain, implying a potential regulation of the KKT23 GNAT domain by the N-terminal region.

**Figure 6.**
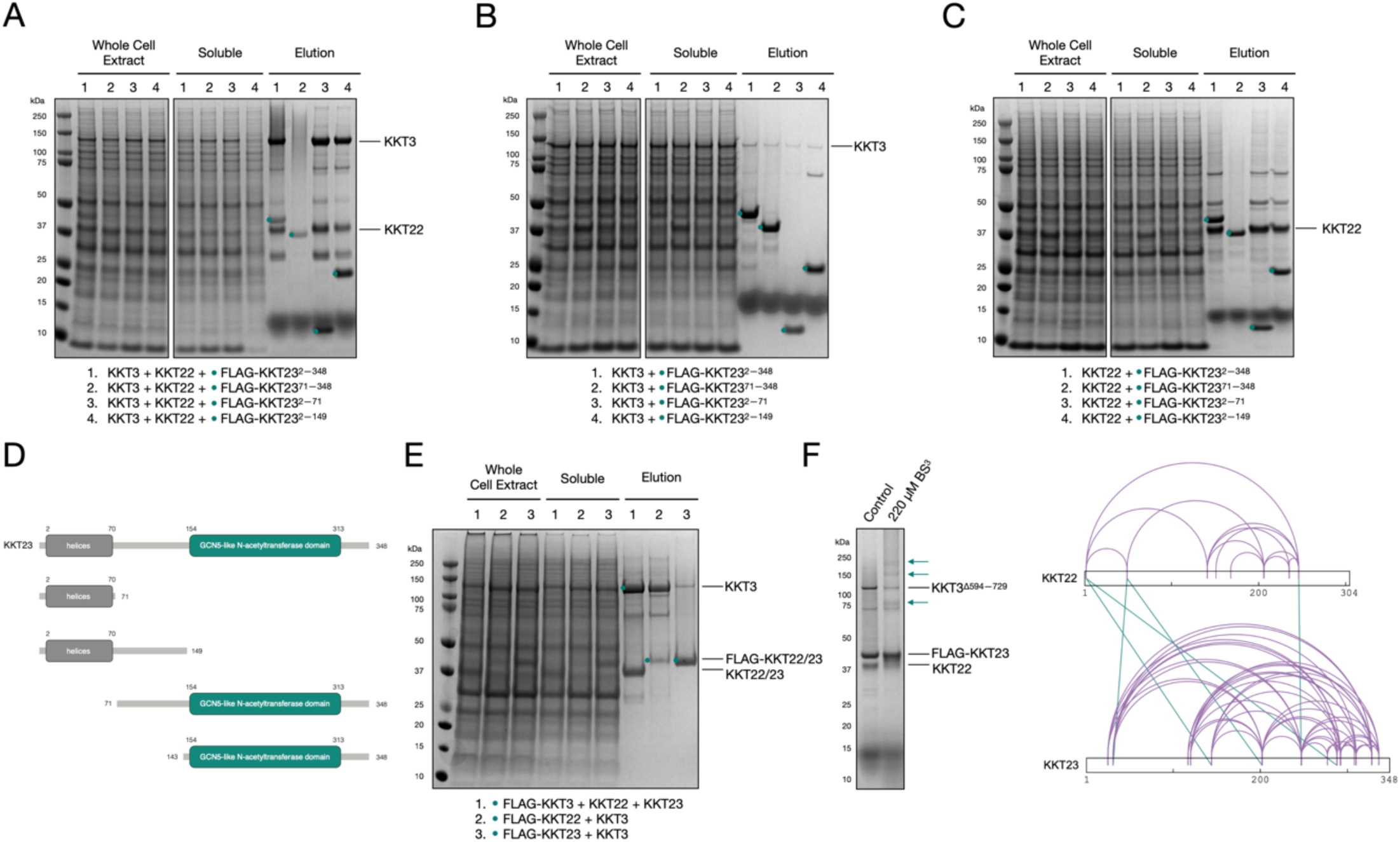
KKT23 forms a complex with KKT3 and KKT22 *in vitro*. A–C. SDS-PAGE showing immunoprecipitation of FLAG-KKT23 fragments with KKT3 and KKT22 (A), KKT3 (B) and KKT22 (C). The migration of FLAG-tagged fragments is labeled with teal dots. FLAG-KKT23 pulls down KKT3 and KKT22 when three proteins are co-expressed. However, KKT23 is not able to pull down KKT3 in the absence of KKT22. KKT23 N-terminal fragment is sufficient to reconstitute the complex composed on three proteins. D. Cartoon schematic depicting boundaries of KKT23 fragments used in (A–C). E. SDS-PAGE showing reconstitution of complexes using full-length KKT3, KKT22 and KKT23 expressed in Sf9 cells. FLAG-KKT3 can pull down both KKT22 and KKT23 (1). Bands corresponding to untagged KKT22 and KKT23 overlap in the gel. The presence of both proteins in the sample was confirmed by mass spectrometry (data not shown). In contrast to FLAG-KKT22 (2), FLAG-KKT23 cannot pull down KKT3 efficiently (3). See also Figure S6. F. SDS-PAGE of the crosslinking reaction using a complex composed of KKT3^Δ594-729^-KKT22-FLAG-KKT23 and 220 µM BS^3^. Due to DNA-contamination of the complex containing the full-length KKT3, deletion mutant of KKT3 was used, where its DNA-binding centromere-localisation domain was removed. The crosslinked species are indicated with teal arrows. Panel on the right illustrates crosslinks detected between KKT22 and KKT23 (green lines). No crosslinks were detected between KKT3 and KKT22 nor KKT23. A complete list of identified crosslinks is shown in Tables S4.

### N-terminal fragment of KKT23 interacts with KKT22

To determine the region of KKT23 that interacts with KKT22, the pull downs were repeated with FLAG-tagged fragments of KKT23. Reconstitution of the three-protein complex could only be observed with the N-terminal fragments of KKT23 (KKT23^2–71^ and KKT23^2–149^) but not with its C-terminal fragment (KKT23^71–348^) (Figure 6A). Like full-length KKT23, the N-terminal fragment of KKT23 co-purified with KKT22 even in the absence of KKT3 (Figure 6C). Based on the observation that the N-terminal region of KKT23 is sufficient to interact with KKT22 in this assay, KKT23^2–150^ was ectopically expressed in *T. brucei* cells and immunoprecipitated to analyse the co-purifying proteins. Indeed, mass spectrometry of the YFP-KKT23^2–150^ immunoprecipitation sample revealed co-purification of many kinetochore proteins, with KKT3 and KKT22 being the top hits (Figure 7A). We next employed a LacO-LacI system [33] to tether the KKT23^2–150^-LacI protein to a non-centromeric locus and found recruitment of KKT22 (Figure 7B). In contrast, no recruitment was observed when KKT23^125–348^-LacI was used (Figure 7B). These results in combination with the immunoprecipitation experiments show the importance of the KKT23 N-terminal region for the interaction with KKT22.

**Figure 7.**
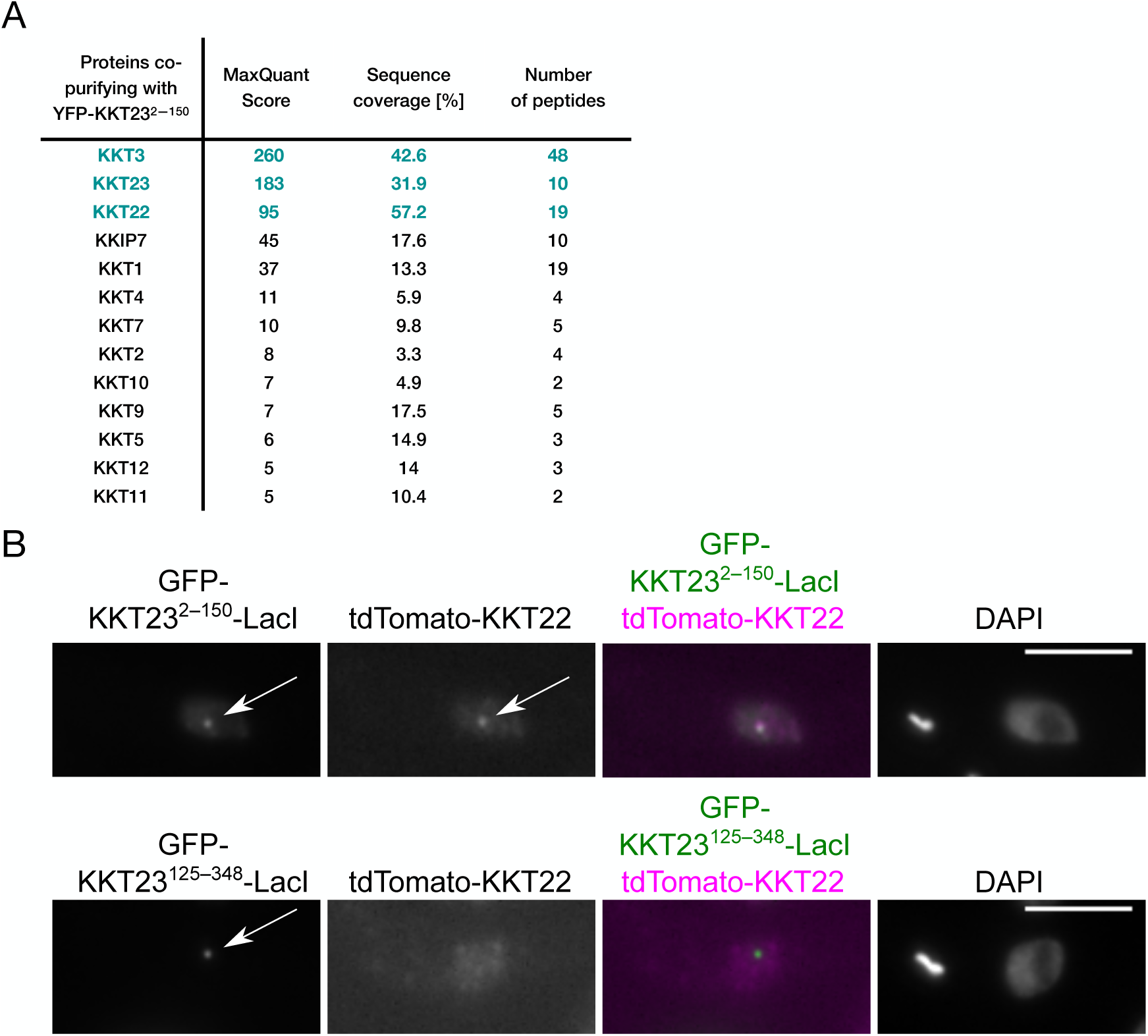
KKT23^2–150^ is sufficient to interact with kinetochore proteins. A. Mass spectrometry analysis of YFP-KKT23^2–150^ immunoprecipitated using anti-GFP antibodies from *T. brucei* cells. The N-terminal fragment of KKT23 co-purifies with a number of kinetochore proteins with KKT3 and KKT22 being the top hits. See Tables S5 for all proteins identified by mass spec. Cell line, BAP2296. B. The N-terminal fragment of KKT23 (KKT23^2–150^, top panel) but not the C-terminal fragment (KKT23^125–348^, lower panel) is sufficient to recruit KKT22 to a non-centromeric locus in *T. brucei* cells (n=5). The GFP-fused proteins expression was induced with 10 ng/ml doxycycline for 24 h. Cell lines, BAP2547 and BAP2546.

### Crystal structure of KKT23^2–70^

Given the importance of the KKT23 N-terminal region, we next aimed to determine its structure using X-ray crystallography. Purification and crystallisation screening of KKT23^2–70^ yielded well diffracting crystals that allowed us to solve the structure at 2.7 Å resolution (Figure 8A, Table 1). The structure comprises three short helices between residues 22–33, 36–49 and 54–62. Due to the lack of electron density in the structure, residues between 2–14 and 65–70 were not modelled. Consequently, we were unable to confirm the secondary structure prediction, which placed an additional helix between residues 5–17 (Figure S1A). To address this issue, we used AlphaFold2 to predict the structure of KKT23^2–70^ (Figure 8B), which indeed revealed an additional helix between residues 5–17 and a longer helix at the C-terminus. Notably, the predicted structure closely matched our crystal structure with an RMSD of 0.9 Å (320 C⍺ atoms). The DALI search of the crystal structure or the AlphaFold2 model did not reveal any obvious structural homologue for KKT23^2–70^ (data not shown). We next used AlphaFold-Multimer and found that the KKT23 N-terminal region is predicted to interact with KKT22 (Figure S6). The helices ⍺1–⍺3 from KKT23 are predicted to interact with ⍺2 and ⍺6 of KKT22 in multiple models with high pLDTT score through hydrophobic and electrostatic interactions.

**Figure 8.**
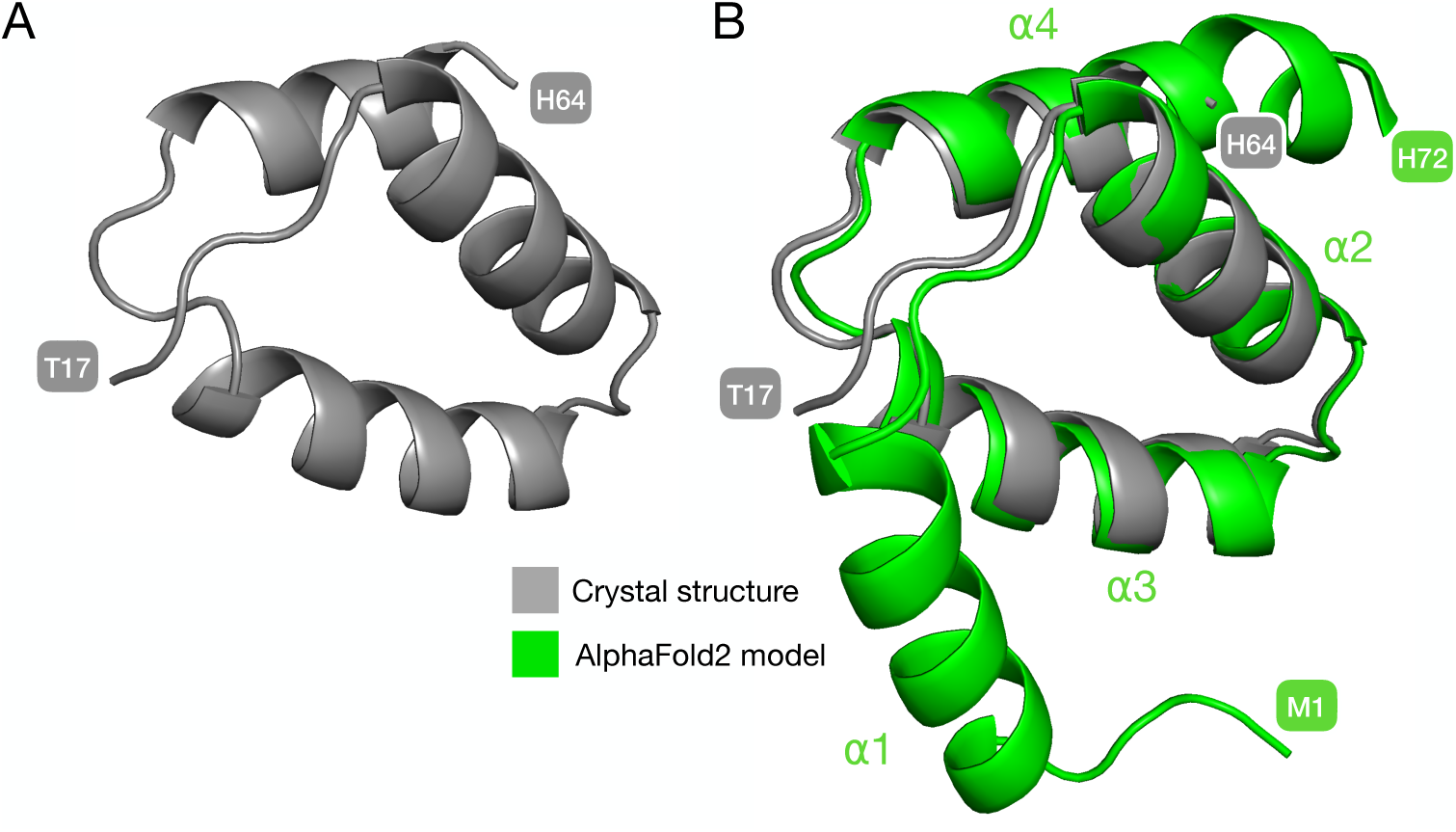
Structure of KKT23^2–70^. A. Ribbon representation of the KKT23^2–70^ structure determined by X-ray crystallography. The model starts at T17 and ends at H64. Residues between 2–16 and 65–70 were not modelled as no electron density was observed for these regions. B. Superposition of the KKT23^2–70^ (grey) with the AlphaFold2 model of KKT23^2–70^ (green). The structures were overlaid using an *align* function in Pymol [13] with an RMSD of 0.9 Å (320 C⍺ atoms). AlphaFold2 predicted an additional N-terminal helix between residues 5–17. No electron density was observed in the crystal structure for this region. Data S1. Ribbon model of the *Trypanosoma brucei* KKT23^125–348^ crystal structure coloured by motifs. Data S2. Ribbon model of the *Trypanosoma brucei* KKT23^125–348^ crystal structure coloured by secondary structure determined with NMR.

### The N-terminal helical domain and the C-terminal GNAT domain are linked by a stretch of unstructured residues

We have determined the structures of KKT23^2–70^ and KKT23^125–348^. However, the region between these two domains, encompassing residues 71–124, is predicted to be disordered (Figure S1B). To gain insight into this region’s structural details, we expressed and purified ^15^N-labeled full-length KKT23. KKT23^2–348^ gives a well-resolved ^1^H-^15^N BEST TROSY spectrum. Many of observed peaks align well with the peaks in the ^1^H-^15^N BEST TROSY spectrum of KKT23^125–348^ (Figure S7), therefore, these peaks were assigned to the GNAT domain. Most of the remaining peaks displayed the higher intensity and narrow dispersion in the ^1^H dimension (Figure S8A), which are often observed for unstructured protein regions. To probe the dynamics of the residues giving rise to these intense peaks, the {^1^H}-^15^N heteronuclear NOE was measured (Figure S8B) [17]. The analysis revealed heteronuclear NOE values below 0.4 for the 39 strongest peaks, indicating that they correspond to unstructured regions of KKT23, most likely the region between residues 71–124. In fact, some of these peaks were assigned to residues in the region of A91 to E100. In conclusion, our data show that the structure of KKT23 consists of an N-terminal helical region, a stretch of unstructured residues, and the GNAT domain at the protein C-terminus.

## Discussion

KKT23 is a unique kinetochore protein that contains a GNAT domain. The fact that KKT23 is conserved in all sequenced kinetoplastids implies that it may play fundamental roles in kinetoplastid kinetochores [6]. Although our previous study showed that KKT23 is essential for cell growth, its structure or function were unknown. Our structural characterisation using X-ray crystallography and NMR confirms that KKT23^125–348^ has an acetyltransferase fold, with the GCN5 histone acetyltransferase being its closest structural homologue. In line with this discovery, we show that KKT23 acetylates the C-terminal tail of histone H2A in *T. brucei.* Although it has been known that the C-terminal tail of histone H2A is hyperacetylated in trypanosomes (K115, K119, K120, K122, K125, K128), the enzyme responsible for them remained unknown [27–31]. Our study provides an answer for this long-standing question by showing that KKT23 contributes to these acetylations. It has been estimated that ∼20% of K115, K119, K120 and 5–10% of K122, K125, K128 is acetylated in proliferating *T. brucei* bloodstream form cells [31]. A previous study performed ChIP-seq using antibodies against the acetylated C-terminal tail (raised against peptides with K115acK120acK122ac or K120acK122acK125ac) and found enrichment at Pol II transcription start regions [34]. Based on the finding that acetylation levels of only K122, K125, and K128 were reduced after KKT23 depletion, it will be important to examine whether acetylation of these specific sites is enriched at the centromeric region. It will also be important to reveal the functional significance of histone H2A acetylation by KKT23. ChIP-seq analysis of KKT3 after depleting KKT23 for 4 days failed to find any obvious change in the position of kinetochore assembly sites (Figure S9). We speculate that KKT23-dependent acetylation of histone H2A could alter chromatin structure directly; acetylation may neutralise lysine’s positive charge and thus destabilise the interaction between the DNA and histones [35, 36]. The C-terminal lysine residues are not ordered in known nucleosome structures, but are likely adjacent to linker DNA based on the presence of H2A C-terminal tail near the entry-exit site of DNA. This could subsequently lead to an enhanced DNA accessibility and transcription [37]. Indeed, transcription has been detected at centromeres in various eukaryotes and is thought to play important roles in ensuring proper kinetochore functions [38]. In fact, small non-coding RNA has been detected for the centromeric repeat sequence of several chromosomes in *T. brucei* [39]. Therefore, KKT23’s acetyltransferase activity may play an important role in ensuring proper kinetochore function by regulating centromeric chromatin. The structural and biochemical insights generated from our analysis of KKT23, a unique kinetochore protein, could open up potential avenues for the rational design of drugs aimed at combating kinetoplastid-associated diseases in the future.

## Acknowledgments

We thank Mark Carrington and Miguel Navarro for sharing reagents. We thank David Staunton for assistance with SEC-MALS and AUC experiments. We thank the Proteomics Core Facility at the EMBL in Heidelberg, especially Mandy Rettel and Jennifer Schwarz, for their support. We thank Krzysztof Kuś for his help with visualisation of crosslinks, Edward Lowe and Gabriele Marcianò for their assistance with determination of crystal structures. We also thank members of Akiyoshi lab for comments and suggestions on the manuscript. Patryk Ludzia was supported by the Boehringer Ingelheim Fonds. Marcus Wilson’s work was supported by the Sir Henry Dale Fellowship form the Wellcome Trust [210493/Z/18/Z] and Medical Research Council (T029471/1). Gauri Deák was supported by BBSRC EASTBIO [BB/M010996/1]. Bungo Akiyoshi was supported by a Wellcome Trust Senior Research Fellowship [210622/Z/18/Z] and a Wellcome Discovery Award [227243/Z/ 23/Z]. The Department of Biochemistry NMR Facility has benefitted from funding provided by the Edward Penley Abraham Fund, the John Fell Fund and the Wellcome Trust. The Lumos mass spectrometer was purchased on a Wellcome Instrument grant [108504]. This work was supported by funding for the Wellcome Discovery Research Platform for Hidden Cell Biology [226791] and we gratefully acknowledge support from the Proteomics core.

## Author contributions

P.L. purified recombinant proteins, solved crystal structures, performed most experiments and data analysis. M.I. performed LacO-LacI tethering and ChIP-seq experiments. G.D and M.W provided reconstituted *T. brucei* and *H. sapiens* nucleosomes. C.S. performed MS analysis. P.L. and C.R. performed and analysed NMR experiments. P.L., C.R., and B.A. designed experiments and wrote the manuscript.

## Rights retention

This research was funded in whole, or in part, by the Wellcome Trust [210622/A/18/Z, 227243/Z/ 23/Z, and 210493/Z/18/Z]. For the purpose of open access, the author has applied a CC BY public copyright licence to any Author Accepted Manuscript version arising from this submission.

## STAR METHODS

### RESOURCE AVAILABILITY

#### Lead contact

Further information and requests for resources and reagents should be directed to and will be fulfilled by the lead contact, Bungo Akiyoshi.

#### Material availability

Plasmids generated in the course of this study can be requested from the Lead contact, Bungo Akiyoshi.

#### Data and code availability

Data generated during this study are included in the manuscript and supplementary information. Protein coordinates have been deposited in the RCSB Protein Data Bank (http://www.rcsb.org/) with accession codes PDB: 9EVQ (native *Trypanosoma brucei* KKT23^125–348^), PDB: 9F5Q (selenomethionine *Trypanosoma brucei* KKT23^125–348^) and PDB: 9EVR (native *Trypanosoma brucei* KKT23^2–70^). The chemical shift assignments for KKT23^125–348^ in the presence of acetyl-CoA have been deposited in the BioMagResBank (http://www.bmrb.wisc.edu) under the accession number 52461. All raw mass spectrometry files and the custom database files used in this study have been deposited to the ProteomeXchange Consortium via the PRIDE partner repository [41, 42] with the dataset identifiers PXD052922 and PXD052923. ChIP-seq data have been uploaded onto the ENA database (PRJEB76406).

### EXPERIMENTAL MODEL AND SUBJECT DETAILS

#### Bacterial culture

Bacterial strains used in this study are listed in Key Resources Table. Bacterial growth conditions can be found in Method Details.

#### Cell culture

*Spodoptera frugiperda* Sf9 cells were cultured in SF-900 SFM media (Gibco) at 27℃ with shaking (160 rpm).

Procyclic form of *Trypanosoma brucei brucei* cells were cultured in SDM-79 medium supplemented with 10% (vol/vol) heat-inactivated fetal calf serum [43].

### METHOD DETAILS

#### Plasmids

All plasmids used in this study are listed in Key Resources Table. KKT fragments used in this study were amplified from *Trypanosoma brucei* genomic DNA and cloned into RSFDuet-1 vector (Novagen) for bacterial expression or into pACEBac2 vector (Invitrogen) for insect cell expression using NEBuilder HiFi DNA Assembly Kit (New England Biolabs). All constructs were sequence verified.

#### Protein expression and purification

The expression of KKT23^125–348^ and KKT23^2–70^ was carried out as follows. Firstly, *E. coli* BL21(DE3) cells were transformed with ∼100 ng of plasmid DNA and then inoculated into 5 ml of 2xTY medium containing 50 µg/ml kanamycin, which was grown overnight at 37°C. For selenomethionine labelling, selenomethionine medium was used instead of 2xTY media (Molecular Dimensions, base mix prepared according to the manufacturer instructions, 5.1 g of nutrients mix and 80 mg of selenomethionine in 1 l of media). In the next morning, each of the 6 l of 2xTY medium with 50 µg/ml of kanamycin was inoculated with 5 ml of the overnight culture and grown at 37°C with shaking (200 rpm) until the OD_600_ reached ∼0.8. Protein expression was induced with 0.2 mM IPTG and was continued for ∼16 hr at 20°C.

Cells were spun down at 3,400 g at 4°C and resuspended in 200 ml of lysis buffer (50 mM sodium phosphate, pH 7.5, 500 mM NaCl, and 10% glycerol) supplemented with protease inhibitors (20 µg/ml leupeptin, 20 µg/ml pepstatin, 20 µg/ml E-64 and 0.4 mM PMSF), benzonase nuclease (500 U/l), and 0.5 mM TCEP. All subsequent steps were performed at 4°C. Bacterial cultures were mechanically disrupted using a French press (1 passage at 20,000 psi) and the soluble fraction was separated by centrifugation at 48,000 g for 30 min. Supernatants were loaded onto 5 ml of TALON beads (Takara Bio) pre-equilibrated with lysis buffer. The beads were then washed with ∼300 ml of the lysis buffer without protease inhibitors and proteins were eluted with 50 mM sodium phosphate pH 7.5, 500 mM NaCl, 10% glycerol, 250 mM imidazole and 0.5 mM TCEP. To cleave off the His-tag, samples were incubated with TEV protease in 1:50 w/w ratio overnight while being buffer-exchanged into 25 mM sodium phosphate, 250 mM NaCl, 5% glycerol, 5 mM imidazole, and 0.5 mM TCEP by dialysis. The samples were then further purified using ion exchange and size exclusion chromatography. To promote binding of protein to the ion exchange column, the samples were diluted with buffer A (25 mM HEPES pH 7.5 and 0.5 mM TCEP) to achieve the final NaCl concentration of 50 mM. Ion exchange chromatography was performed using a 6 ml RESOURCE S column for KKT23^125–348^ and a RESOURCE Q for KKT23^2–348^ and KKT23^2–70^, both pre-equilibrated with 5% of buffer B (25 mM HEPES pH 7.5, 1 M NaCl and 0.5 mM TCEP). Protein was eluted with a linear gradient from 5% to 100% of buffer B, concentrated using 10-kD MW Amicon concentrators (Millipore), and loaded on Superdex 75 16/60 (GE Healthcare) columns to further purify and buffer exchange into 25 mM HEPES pH 7.5, 150 mM NaCl with 0.5 mM TCEP. Fractions containing KKT23 were pooled, concentrated and flash-frozen in liquid nitrogen for -80°C storage.

#### Expression of isotopically labeled KKT23 fragments

The E. coli BL21(DE3) cells were transformed with ∼100 µg of plasmid DNA and were inoculated into 40 ml of 2xTY media containing 50 µg/ml kanamycin. Cells were grown at 37°C overnight. The next day, cells were spun down at 3,400 g for 10 min and resuspended in 40 ml of M9 minimal medium containing 50 µg/ml kanamycin, 1g/l ^15^NH_4_Cl and 4g/l [^13^C]-D-glucose (CIL) as the sole nitrogen and carbon sources. Next, the resuspended culture was inoculated into 1 l of M9 minimal medium supplemented with 1g/l ^15^NH_4_Cl, 4g/l [^13^C]-D-glucose and 50 µg/ ml kanamycin. Cells were grown at 37°C to an OD_600_ of 0.9–1.0 and the protein expression was induced using 0.4 mM IPTG. The protein expression was continued overnight at 20°C with shaking (200 rpm).

#### Expression and purification from insect cells

Typically, 50 ml of Sf9 cells at 1–1.2 million cells/ml were infected with 0.5 ml of P3 baculovirus for 72h before harvesting. Cells were pelleted at 700 g for 10 min and resuspended in 2 ml of BH0.25 (25 mM HEPES, pH 8.0, 0.2 % NP-40, 2 mM MgCl_2_, 0.1 mM EDTA, 0.5 mM EGTA, 10% glycerol, and 250 mM NaCl) supplemented with protease inhibitors (20 mg/ml leupeptin, 20 mg/ml pepstatin, 20 mg/ml E-64, and 0.4 mM PMSF) and 25U of benzonase nuclease. Cells were homogenised on ice by sonication (three rounds of 15 s pulse and 1 min pause) followed by centrifugation for 30 min at 45 000 g. The supernatant was incubated with pre-equilibrated 0.1 ml of anti-FLAG M2 affinity gel (Sigma) for 2.5 h with constant rotation at 4℃. This was followed by five washes with BH0.25 supplemented with 0.5 mM TCEP (1 ml each). To elute the protein, the beads were incubated with 0.1 ml of BH0.25 containing 0.5 mg/ml 3xFLAG peptide (Sigma) for 30 min at RT with gentle agitation. Supernatant containing the eluted protein was separated from the beads using centrifugation and flash frozen in liquid nitrogen for the -70℃ storage.

#### Size Exclusion Chromatography with Multi-Angle Light Scattering

MALS experiments were performed during size exclusion chromatography on analytical column SD75 HR10/300 equilibrated with 25 mM HEPES pH 7.5, 150 mM NaCl and 0.5 mM TCEP. Protein elution was monitored via online static light-scattering (DAWN HELEOS 8+, Wyatt Technology), differential refractive index (Optilab T-rEX, Wyatt Technology) and UV (SPD-20A, Shimadzu) detectors. Data were analysed and molecular weight calculated using the ASTRA software package (Wyatt Technology). KKT23^2–348^ was loaded on size exclusion column at 70 µM in the presence of 140 µM acetyl coenzyme A.

#### Analytical Ultracentrifugation

The analytical ultracentrifugation experiments were carried out using a Beckman An-60 Ti rotor in a Beckman Optima XL-1 analytical ultracentrifuge at 20°C. KKT23^2–348^ sample was prepared at 70 µM in the presence of 140 µM acetyl coenzyme A in 25 mM HEPES pH 7.5, 150 mM NaCl and 0.5 mM TCEP. Scans were recorded using interference optics. Multiple scans were fitted to a continuous size distribution using SEDFIT [44]. The buffer density and viscosity were calculated with SEDNTERP [45]. The sample partial specific volumes were estimated from the amino acid sequences, also using SEDNTERP.

#### Crystallisation

All crystals were obtained in sitting drop vapour diffusion experiments in 96-well plates, using drops of 200 nl overall volume, mixing protein and mother liquor in a 1:1 ratio.

Crystals of *Trypanosoma brucei* KKT23^2–70^ (12.0 mg/ml) were grown at 18℃ in ProPlex HT-96 E7 solution (Molecular Dimensions) containing 0.1M Sodium phosphate, pH 6.5, 12% w/ v PEG 8000. Crystals appeared after 40 days. Crystals were briefly transferred into mother liquor prepared with addition of 25% glycerol prior to flash-cooling by plunging into liquid nitrogen.

For the native and selenomethionine KKT23^125–348^ samples, before setting the trays, they were supplemented with 1.6 and 0.98 mM acetyl-CoA, respectively. Native crystals of *Trypanosoma brucei* KKT23^125–348^ (14.5 mg/ml) were grown at 4℃ in MIDAS HT-96 B9 solution (Molecular Dimensions) containing 0.1 M sodium tartrate dibasic dehydrate, 0.1 M HEPES pH 7.0, 20% SOKOLAN PA 25 CL. Selenomethionine crystals of *Trypanosoma brucei* KKT23^125–348^ (12.5 mg/ml) were grown at 4℃ in MIDAS HT-96 F7 solution (Molecular Dimensions) containing 20% v/v jeffamine M-2070, 20% v/v DMSO. Crystals were briefly transferred into mother liquor prepared with addition of 25% glycerol prior to flash-cooling by plunging into liquid nitrogen.

#### Diffraction Data Collection and Structure Determination

X-ray diffraction data from *Trypanosoma brucei* KKT23^2–70^ were collected at the I04 beamline at the Diamond Light Source (Harwell, UK). The structure was solved using *ab initio* macromolecular phasing software, ARCIMBOLDO LITE [46] followed by initial model building with BUCCANEER [47]. Further manual model building and refinement were completed iteratively using COOT [48] and PHENIX [49]. The data sets used for the final refinement were scaled to the high-resolution limit of 2.7 Å.

X-ray diffraction data from selenomethionine crystals of *Trypanosoma brucei* KKT23^125– 348^ were collected at the I03 beamline at the Diamond Light Source (Harwell, UK). The structure was solved using CRANK2 software [50] followed by initial model building with BUCCANEER [47]. Further manual model building and refinement were completed iteratively using COOT [48] and PHENIX [49].

X-ray diffraction data from native crystals of *Trypanosoma brucei* KKT23^125–348^ were collected at the I03 beamline at the Diamond Light Source (Harwell, UK). The structure was solved using PHASER [51] and selenomethionine structure as a model. This was followed by initial model building with BUCCANEER [47]. Further manual model building and refinement were completed iteratively using COOT [48] and PHENIX [52]. The dataset used for the final refinement were scaled to the high-resolution limit of 1.75 Å followed by two final rounds of refinement using PHENIX.

#### NMR spectroscopy and analysis on NMR data

NMR samples used throughout this study were prepared in 25 mM HEPES pH 7.3, 150 mM NaCl, 0.5 mM TCEP and 95% H_2_O/5% D_2_O. KKT23^2–348^ and KKT23^125–348^ were additionally supplemented with two-fold excess of acetyl coenzyme A. All NMR spectra were acquired using a 750 MHz spectrometer equipped with a Bruker Avance III HD console and a 5 mm TCI CryoProbe. All NMR data were processed using NMRPipe [53] and analysed using CCPN Analysis [54]. ^1^H, ^13^C and ^15^N chemical shifts of KKT23^125–348^ were analysed using TALOS-N [16] to predict secondary structure propensities. The {^1^H}-^15^N heteronuclear NOE was measured for KKT23^2–348^ (0.25 mM + 0.5 mM acetyl-CoA) and KKT23^125–348^ (0.35 mM + 0.7 mM acetyl-CoA) using the TROSY-based heteronuclear NOE experiment recorded with and without ^1^H saturation for 4.0 sec at 750 MHz [55]. The data sets were acquired using 100 complex t1 increments, 88 (KKT23^125–348^) or 116 (KKT23^2–348^) scans per increment and with a ^15^N sweep width of 2283.10 Hz. 1K complex data points were recorded in the F2 dimension with a sweep width of 10638.30 Hz. Data were collected at 20℃ for KKT23^2–348^ and KKT23^125–348^. The {^1^H}-^15^N NOE was calculated as the ratio of the peak intensities in the spectra recorded with and without ^1^H saturation. Peak heights were determined using CCPN Analysis [54]. Uncertainties in the {^1^H}-^15^N NOE values were estimated from 500 Monte Carlo simulations using the baseline noise as a measure of the error in the peak heights.

#### Reconstitution of *T. brucei* and *H. sapiens* nucleosome core particles

Expression and purification of histones, and reconstitution of nucleosome core particles was done as previously described [26, 56]. Briefly, histones were expressed in E. coli BL21(DE3)RIL cell inclusion bodies. The inclusion bodies were lysed, sonicated, washed, unfolded, and dialysed into 7 M urea, 15 mM Tris pH 7.5, 100 mM NaCl, 1 mM EDTA, and 5 mM BME. The lysates were then purified by cation exchange chromatography using a 20 column volume elution gradient of 0–80% 7 M urea, 15 mM Tris pH 7.5, 1 M NaCl, 1 mM EDTA, and 5 mM BME. Protein containing fractions were pooled, dialysed into 2 mM BME, lyophilised, and stored at -20°C.

Histone octamers were assembled by resuspending the purified, lyophilised histones in 7 M guanidine-HCl, 20 mM Tris pH 7.5, and 10 mM DTT, mixing them at a 1.2:1.2:1:1 (H2A:H2B:H3:H4) molar ratio, and dialysing the mixture into 15 mM Tris pH 7.5, 2 M NaCl, 1 mM EDTA, and 5 mM BME. The resulting complex was purified by size exclusion chromatography (Superdex S200).

Widom 601 145 bp DNA for histone octamer wrapping was isolated from a pUC57 plasmid encoding 8 copies of the 145 bp DNA with EcoRV overhangs using EcoRV restriction enzyme digestion, PEG precipitation, and ethanol precipitation.

Nucleosome core particles were then reconstituted by mixing either *T. brucei* or *H. sapiens* histone octamers with the Widom 601 145 bp DNA via a salt dialysis gradient. The nucleosome core particles were subsequently dialysed into a final storage buffer (15 mM HEPES pH 7.5, 25 mM NaCl, 2.5% glycerol, and 1 mM DTT).

#### *In vitro* histone acetyltransferase assays on reconstituted nucleosomes

Prior to the assay, KKT23^2–348^ was buffer exchanged into the HAT buffer (50 mM Tris-HCl pH 8.0, 50 mM NaCl, 5% glycerol, 1 mM DTT, 0.1 mM EDTA and 10 mM sodium butyrate) using a gel filtration column SD75 16/600. The eluted protein was diluted to 2 µM and 60 µM stocks and flash-frozen in liquid nitrogen for storage at -70℃. Prior to mixing the enzyme with reconstituted *T. brucei* and human nucleosomes [26], acetyl-CoA sodium salt (Sigma Aldrich) was added to the enzyme up to a concentration of 200 µM. The final concentration of the enzyme and substrates was 30 µM and 1 µM, respectively. Then, the enzyme was mixed with the substrate in 1:1 v/v ratio and incubated at 30°C for 1 h. Following incubation, samples were boiled for 3 min and resolved using SDS-PAGE electrophoresis, followed by mass spectrometry (see below) or Western blot analysis. To access composition and quantify loading, samples were resolved using NuPage 4–12% gradient gels (Invitrogen) and stained with SimplyBlue (Invitrogen). For the Western blot analysis, proteins from SDS-PAGE gels were transferred onto nitrocellulose membranes, and blocking was done with 5% skimmed milk in PBST. The membranes were then incubated with primary anti pan-acetyl lysine antibodies (#9814S from Cell Signaling) at 1:1000 v/v dilution in PBST overnight at 4℃. Following incubation, the membrane was washed three times with PBST and incubated with anti-rabbit horseradish peroxidase-linked secondary antibodies for 1h at RT (at 1:1000 v/v dilution in PBST). Acetylated proteins were detected by chemiluminescence using SuperSignal West Pico PLUS Chemiluminescent Substrate (Thermo Scientific).

#### Trypanosome cells

Trypanosome cell lines were derived from *T. brucei* SmOxP927 procyclic form cells (TREU 927/4 expressing T7 RNA polymerase and the tetracycline repressor to allow inducible expression; [57]. *Trypanosoma brucei* cell cultures were grown at 28°C in SDM-79 medium supplemented with 10% (vol/vol) heat-inactivated fetal calf serum [43]. LacO-LacI tethering experiments were carried out as described previously using the LacO array inserted at the rDNA locus [33, 58], and expression of GFP-NLS-LacI fusions was induced by the addition of 10 ng/ml doxycycline for 24 hr. ChIP-seq analysis was performed as described previously [4, 59].

#### Microscopy

Trypanosome cells expressing GFP-NLS-LacI fusion with KKT23^2–150^ or KKT23^128–348^ were fixed with formaldehyde as previously described [5]. Images were captured at room temperature on a Zeiss Axioimager.Z2 microscope (Zeiss) installed with ZEN using a Hamamatsu ORCA-Flash4.0 camera with 63x objective lenses (1.40 NA). Typically, 15 optical slices spaced 0.2 µm apart were collected. Images were analysed in ImageJ/Fiji [60].

#### Immunoprecipitation of histones and KKT23^2–150^ from trypanosomes

Initially, a 40 ml culture of procyclic cells expressing either H2B-GFP (BAP2596) or GFP-KKT23^2–150^ (BAP2296) was prepared at a concentration of 1.2 x 10^7^ cells/ml. Subsequently, these cells were inoculated into a larger culture volume of 400 ml at a concentration of 3 x 10^5^ cells/ml, evenly distributed among three flasks. After 24h, KKT23 RNAi-mediated knockdown was induced with doxycycline at 1 µg/ml final concentration in BAP2596. Expression of KKT23^2–150^ in BAP2296 was induced with 10 ng/ml doxycycline. The flasks were then incubated for 1 more day at 27°C until the cells reached approximately 1 x 10^7^ cells/ml. After the incubation period, cells were centrifuged at 800 g for 15 min at room temperature. The supernatant was discarded, and cells were washed with 40 ml of PBS. Following centrifugation at 800 g for 5 min, cells were resuspended in 25 ml of PEME buffer (100 mM PIPES-NaOH pH 6.9, 2 mM EGTA, 1 mM MgSO_4_, and 0.1 mM EDTA) containing 1% NP-40 and protease inhibitors (Leupeptin, Pepstatin, E-64, 10 µg/ml each, and 0.2 mM PMSF), phosphatase inhibitors (1 mM sodium pyrophosphate, 2 mM Na-beta-glycerophosphate, 0.1 mM Na_3_VO_4_, 5 mM NaF, and 100 nM microcystin-LR), and a histone deacetyltransferase inhibitor (50 mM sodium butyrate). After the 5 min incubation, the lysed cells were centrifuged at 1800 g for 15 min at room temperature. The supernatant was discarded, and the pellet fraction containing the cytoskeleton was transferred on ice and resuspended in 1.5 ml of BH0.15 buffer (25 mM HEPES pH 8.0, 2 mM MgCl_2_, 0.1 mM EDTA pH 8.0, 0.5 mM EGTA pH 8.0, 1% NP-40, 150 mM KCl, and 15% glycerol) supplemented with protease inhibitors (Leupeptin, Pepstatin, E-64, 10 µg/ml each, and 0.2 mM PMSF), phosphatase inhibitors (1 mM sodium pyrophosphate, 2 mM Na-beta-glycerophosphate, 0.1 mM Na_3_VO_4_, 5 mM NaF, and 100 nM microcystin-LR), and a histone deacetyltransferase inhibitor (50 mM sodium butyrate). Samples were then subjected to sonication to solubilise proteins (50% amplitude, 12 s pulse, 60 s pause, 3 cycles). Next, the samples were incubated with 60 µl of Protein-G magnetic beads (Invitrogen) slurry, preconjugated with 12 µg of mouse monoclonal anti-GFP antibodies (Roche, 11814460001) using dimethyl pimelimidate. Following the 2.5 h incubation with constant rotation, the samples were washed with BH0.15 buffer (5 x 1 ml) and eluted with 60 µl of elution buffer (50 mM Tris pH 8.3, 1 mM EDTA and 0.3% SDS) with gentle agitation for 25 min at room temperature. Finally, the samples were frozen and stored at -70°C until further analysis by mass spectrometry.

#### Mass spectrometry: In-solution trypsin digest for immunoprecipitates of H2B-GFP and GFP-KKT23^2–150^ purified from trypanosomes

Reduction of disulphide bridges in cysteine-containing proteins was performed with dithiothreitol (56°C, 30 min, 10 mM in 50 mM HEPES, pH 8.5). Reduced cysteines were alkylated with 2-chloroacetamide (room temperature, in the dark, 30 min, 20 mM in 50 mM HEPES, pH 8.5). Samples were prepared using the SP3 protocol (Hughes et al., 2019) and trypsin (sequencing grade, Promega) was added in an enzyme to protein ratio 1:50 for overnight digestion at 37°C. Next day, peptide recovery in HEPES buffer by collecting supernatant on magnet and combining with second elution wash of beads with HEPES buffer. For further sample clean up an OASIS HLB µElution Plate (Waters) was used. The samples were dissolved in 10 µL of reconstitution buffer (96:4 water: acetonitrile, 1% formic acid and analysed by LC-MS/MS using a QExactive (GFP-KKT23^2–150^ IP sample) at the proteomics core facility at EMBL Heidelberg, Orbitrap Fusion Lumos mass spectrometer (Thermo) at EMBL Heidelberg (2 replicates of H2B-GFP IP samples), or Orbitrap Fusion Lumos mass spectrometer the Proteomics Facility at Wellcome Centre for Cell Biology (1 replicate of H2B-GFP IP samples) as follows.

##### Mass spectrometry data acquisition on QExactive at EMBL (for GFP-KKT23^2–150^ IP)

An UltiMate 3000 RSLC nano LC system (Dionex) fitted with a trapping cartridge (µ-Precolumn C18 PepMap 100, 5µm, 300 µm i.d. x 5 mm, 100 Å) and an analytical column (nanoEase™ M/Z HSS T3 column 75 µm x 250 mm C18, 1.8 µm, 100 Å, Waters). Trapping was carried out with a constant flow of trapping solution (0.05% trifluoroacetic acid in water) at 30 µL/min onto the trapping column for 6 minutes. Subsequently, peptides were eluted via the analytical column running solvent A (0.1% formic acid in water) with a constant flow of 0.3 µL/min, with increasing percentage of solvent B (0.1% formic acid in acetonitrile). The outlet of the analytical column was coupled directly to an Orbitrap QExactive plus Mass Spectrometer (Thermo) using the Nanospray Flex™ ion source in positive ion mode. The peptides were introduced into the QExactive plus via a Pico-Tip Emitter 360 µm OD x 20 µm ID; 10 µm tip (New Objective) and an applied spray voltage of 2.2 kV. The capillary temperature was set at 275°C. Full mass scan was acquired with a mass range 350-1500 m/z in profile mode with resolution of 70000. The filling time was set at a maximum of 50 ms with a limitation of 3×10^6^ ions. Data dependent acquisition (DDA) was performed with the resolution of the Orbitrap set to 17500, with a fill time of 120 ms and a limitation of 5×10^4^ ions. A normalised collision energy of 30 was applied. Dynamic exclusion time of 30 s was used. The peptide match algorithm was set to ‘preferred’ and charge exclusion ‘unassigned’, charge states 1 and 2 were excluded. MS^2^ data was acquired in centroid mode.

##### Mass spectrometry data acquisition on Lumos at EMBL (for 2 replicates of H2B IP experiments)

An UltiMate 3000 RSLC nano LC system (Dionex) equipped with an µ-Precolumn (C18 PepMap 100, 5µm, 300 µm i.d. x 5 mm, 100 Å) and an analytical column (nanoEase M/Z HSS T3 column 75 µm x 250 mm C18, 1.8 µm, 100 Å, Waters) was coupled directly to an Orbitrap Fusion Lumos Tribrid Mass Spectrometer (Thermo) using the Nanospray Flex ion source in positive ion mode. Sample trapping was carried out with a constant flow of 0.05% trifluoroacetic acid in water at 30 µL/min onto the pre-column for 4 minutes. Subsequently, peptides were eluted via the analytical column running solvent A (0.1% formic acid in water, 3% DMSO) with a constant flow of 0.3 µL/min, with increasing percentage of solvent B (0.1% formic acid in acetonitrile, 3% DMSO) from 2% to 8% in 6 min, to 25% in 41 min, to 40% in 5 min to 80% in 4 min and a re-equilibration back to 2% B for 4 min. The peptides were introduced into the Fusion Lumos via a Pico-Tip Emitter 360 µm OD x 20 µm ID; 10 µm tip (CoAnn Technologies) and an applied spray voltage of 2.4 kV. The capillary temperature was set at 275°C. Full mass scan (MS1) was acquired with a mass range 375-1750 m/z in profile mode in the orbitrap with resolution of 120000. The filling time was set at maximum of 50 ms, the AGC target to standard. Data dependent acquisition (DDA) was performed with the resolution of the Orbitrap set to 15000, with a fill time of 54 ms and an AGC target of 400%. A normalised collision energy of 34 was applied. MS2 data was acquired in profile mode.

##### Mass spectrometry data acquisition on Lumos at Edinburgh (for 1 replicate of H2B IP experiments)

Following digestion, samples were cleaned and peptides were concentrated as described by [61]. Peptides were then eluted in 40 µL of 80% acetonitrile in 0.1% TFA and concentrated down to 1 µL by vacuum centrifugation (Concentrator 5301, Eppendorf, UK). The peptide sample was then prepared for LC-MS/MS analysis by diluting it to 6 µL by 0.1% TFA. LC-MS analyses were performed on an Orbitrap Fusion Lumos Tribrid Mass Spectrometer (Thermo Fisher Scientific, UK) both coupled on-line, to an Ultimate 3000 HPLC (Dionex, Thermo Fisher Scientific, UK). Peptides were separated on a 50 cm (2 µm particle size) EASY-Spray column (Thermo Scientific, UK), which was assembled on an EASY-Spray source (Thermo Scientific, UK) and operated constantly at 50°C. Mobile phase A consisted of 0.1% formic acid in LC-MS grade water and mobile phase B consisted of 80% acetonitrile and 0.1% formic acid. Peptides were loaded onto the column at a flow rate of 0.3 µL min-1 and eluted at a flow rate of 0.25 µL min-1 according to the following gradient: 2 to 40% mobile phase B in 150 min and then to 95% in 11 min. Mobile phase B was retained at 95% for 5 min and returned back to 2% a minute after until the end of the run (190 min). MS1 scans were recorded at 120,000 resolution (scan range 350-1500 m/z) with an ion target of 4×10^5^, and injection time of 50ms. MS2 was performed in the ion trap at a rapid scan mode, with ion target of 2×10^4^ and HCD fragmentation [62] with normalised collision energy of 27. The isolation window in the quadrupole was 1.4 Thomson. Only ions with charge between 2 and 6 were selected for MS2. Dynamic exclusion was set at 60 s.

Peptides were identified by searching tandem mass spectrometry spectra against the *T. brucei* protein database with MaxQuant (version 2.0.1) with carbamidomethyl cysteine set as a fixed modification and oxidisation (Met), acetylation (Lys),and phosphorylation (Ser and Thr) set as variable modifications. For the analysis of GFP-KKT23^2–150^ sample, up to two missed cleavages were allowed. For the analysis of H2B-GFP samples, up to six missed cleavages were allowed. The first peptide tolerance was set to 10 ppm. To compare acetylation levels on histone H2A in WT and KKT23-depleted cells, intensities of acetylated peptides were normalised by the total amount of histone H2A protein detected in each sample.

#### Mass Spectrometry: In-gel digestion for nucleosomes acetylated using recombinant KKT23

The bands corresponding to H2A/H3, H4, or H2B were cut from the gel and subjected to in-gel digestion with trypsin (sequencing grade, Promega). Peptides were extracted from the gel pieces by sonication for 15 minutes, followed by centrifugation and supernatant collection. A solution of 50:50 water: acetonitrile, 1% formic acid (2 x the volume of the gel pieces) was added for a second extraction and the samples were again sonicated for 15 minutes, centrifuged and the supernatant pooled with the first extract. The pooled supernatants processed using speed vacuum centrifugation. The samples were dissolved in 10 µL of reconstitution buffer (96:4 water: acetonitrile, 1% formic acid and analysed by LC-MS/MS using an Orbitrap Fusion Lumos mass spectrometer (Thermo) at the proteomics core facility at EMBL Heidelberg as described above.

Peptides were identified by searching tandem mass spectrometry spectra against the *T. brucei* protein database with MaxQuant (version 2.0.1) with carbamidomethyl cysteine set as a fixed modification, while oxidisation (Met) and acetylation (Lys) were set as variable modifications. Up to six missed cleavages were allowed. Total peptide sequence coverages were as follows: 99.3% for H2A, 99.1% for H2B, 99.2% for H3, 99.0% for H4. The first peptide tolerance was set to 10 ppm.

#### Chemical crosslinking mass spectrometry (XL-MS) of KKT23 complex

Prior to the experiment, bis(sulfosuccinimidyl)suberate, BS^3^ (Thermo Fisher) was equilibrated at room temperature for 2 h and then resuspended to 2.2 mM in distilled water. Immediately after, 2 µl of the crosslinker were mixed with 18 µl of ∼3 µM protein complex in 25 mM HEPES pH 8.0, 250 mM NaCl, 10% glycerol, 2 mM MgCl_2_, 0.1 mM EDTA, 0.5 mM EGTA-KOH, 0.1% NP40 and 0.5 mg/ ml FLAG peptide resulting final BS^3^ concentration of 220 µM. The crosslinking reaction was incubated on ice for 60 min. To quench the reactions, Tris-HCl at pH 8.0 was added up to 50 mM concentration and the samples were incubated for 10 min at RT. Next, the samples were fresh-frozen in liquid nitrogen and stored at -70℃ until mass spectrometry analysis. To estimate the efficiency of the crosslinking reactions, samples were resolved on Nu-PAGE 4–12% gradient polyacrylamide gel (Invitrogen), which was stained using SimplyBlue (Invitrogen). In-solution trypsin digestion with SP3 was carried out as described above at the proteomics core facility at EMBL Heidelberg.

An UltiMate 3000 RSLC nano LC system (Dionex) equipped with an µ-Precolumn (C18 PepMap 100, 5µm, 300 µm i.d. x 5 mm, 100 Å) and an analytical column (nanoEase M/Z HSS T3 column 75 µm x 250 mm C18, 1.8 µm, 100 Å, Waters) was coupled directly to an Orbitrap Fusion Lumos Tribrid Mass Spectrometer (Thermo) using the Nanospray Flex ion source in positive ion mode. Trapping was carried out with a constant flow of 0.05% trifluoroacetic acid in water at 30 µL/ min onto the pre-column for 6 minutes. Subsequently, peptides were eluted via the analytical column running solvent A (0.1% formic acid in water, 3% DMSO) with a constant flow of 0.3 µL/ min, with increasing percentage of solvent B (0.1% formic acid in acetonitrile, 3% DMSO) from 2% to 6% in 2 min, to 8% in 2 min, to 30% in 41 min to 40%, in 5 min, to 85% in 7 min, followed by a re-equilibration back to 2% B in 3 min. The peptides were introduced into the Orbitrap Fusion Lumos via a Pico-Tip Emitter 360 µm OD x 20 µm ID; 10 µm tip (New Objective) and an applied spray voltage of 2.4 kV, instrument was operated in positive mode. The capillary temperature was set at 275°C. Full mass scans were acquired for a mass range 350-1700 m/z in profile mode in the orbitrap with resolution of 120000. The filling time was set to auto and the AGC target was set to standard. The instrument was operated in data dependent acquisition (DDA) mode and MSMS scans were acquired in the Orbitrap (Resolution at 30000), with a fill time of up to 100 ms and AGC target of 400%. A normalised collision energy of 32 was applied. MS2 data was acquired in profile mode. RAW MS files were searched by the pLink2 software [63], with carbamidomethyl cysteine set as a fixed and oxidisation (Met) set as variable modifications. Up to two missed cleavages were allowed. Precursor tolerance was set to 10 ppm. All the identified cross-links are shown in Table S4 (FDR 5%). Cross-links were plotted using xiView [64]. Only crosslinks with a score better than E^-1.5^ are shown in Figure 6.

#### Multiple Sequence Alignment

The DNA and protein sequences for KKT23 were retrieved from the TriTryp database [65], UniProt [66] or a published study [67]. Searches for the homologous proteins were done using BLAST in the TriTryp database [65] or using hmmsearch using manually prepared hmm profiles (HMMER, version 3.0) [68]. The multiple sequence alignments were performed with MAFFT (L-INS-i method, version 7) [69] and visualised with the Clustalx colouring scheme in Jalview (version 2.11) [70].

### *In silico* structure and interaction predictions

Structures and interactions were predicted with AlphaFold-Multimer-v2.3.1 [71, 72] through ColabFold version 1.5.5 using MMseqs2 (UniRef+Environmental) and default settings [73]. The following command was used to map pLDDT score onto the AF2 predicted structure models: spectrum b, rainbow_rev, maximum = 100, minimum = 50.

---

**Figure S1.**
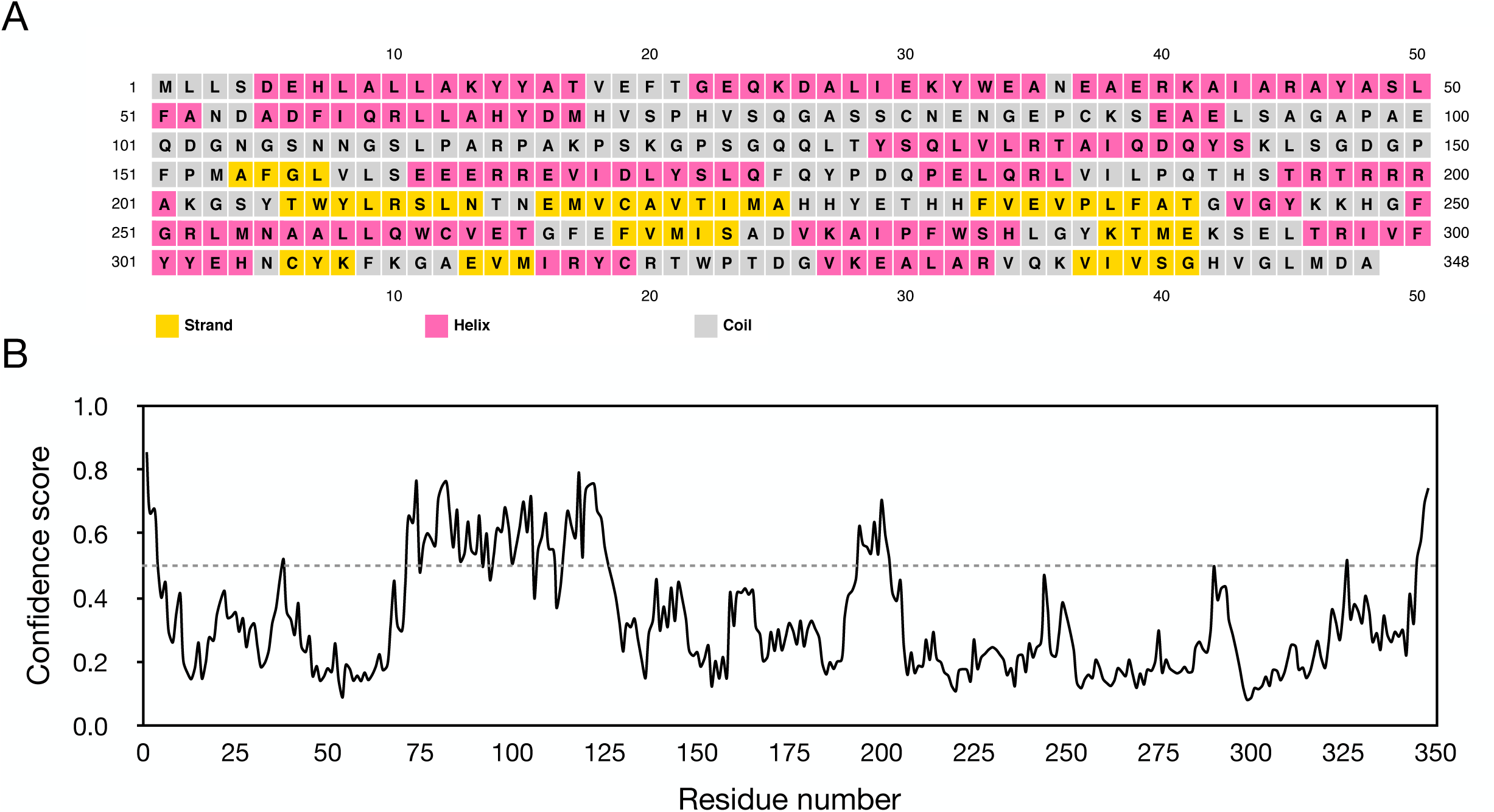
Analysis of the KKT23 primary sequence. A. The secondary structure of KKT23 was analysed using PSIPRED (McGuffin et al., 2000). Each box represents an amino acid labelled with one letter code and coloured according to the predicted secondary structure (yellow for β-strand, pink for ⍺-helix and grey for coil). B. The disorder probability is plotted against the KKT23 sequence. The prediction was made using the DisEMBL server (Linding et al., 2003). The dashed line represents the threshold (0.5) between order and disorder. KKT23 is mainly ordered excluding the regions between 72—126 and 194—202 that have disorder probability values >0.5. This is consistent with the secondary structure prediction shown in (A).

**Figure S2.**
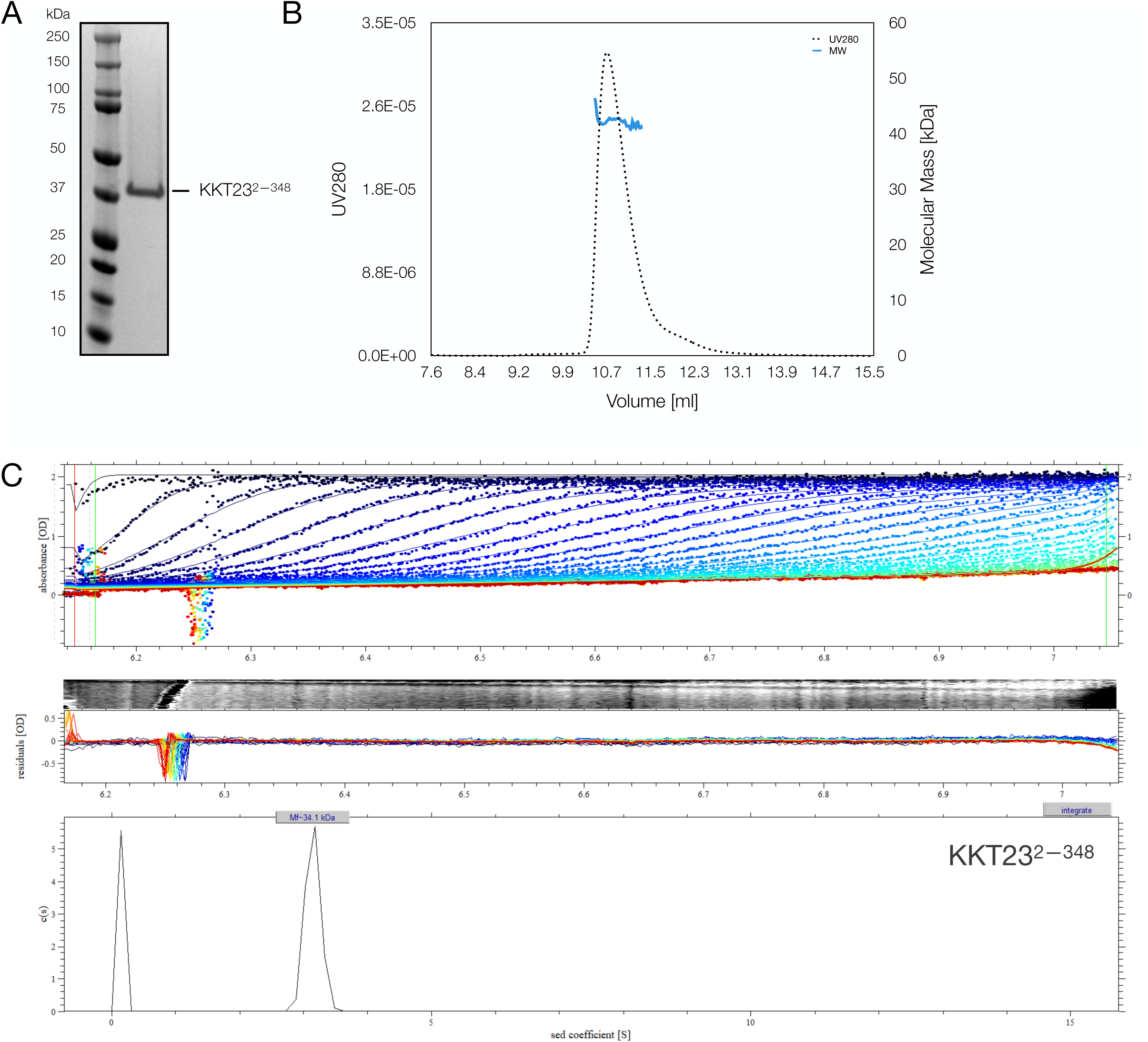
KKT23 is a monomer in solution. A. SDS-PAGE gel stained with coomassie brilliant blue of the full-length recombinant KKT23. The gel depicts a fraction from size exclusion chromatography run on SD75 16/60 column. B. SEC-MALS elution profile of 50 μM KKT23^2—348^ in the presence of 100 μM acetyl-CoA. The UV_280_ signal (dotted line) is plotted against the elution volume, while the molecular weight is indicated as a thick line. The estimated molecular weight (∼40 kDa) matches the calculated value based on the sequence (39 kDa), suggesting that KKT23 is a monomer. C. AUC analysis of 50 μM KKT23^2—348^ in the presence of 100 μM acetyl-CoA. The AUC results are consistent with the SEC-MALS analysis in (A), showing that KKT23 sediments as a 34 kDa monomer.

**Figure S3.**
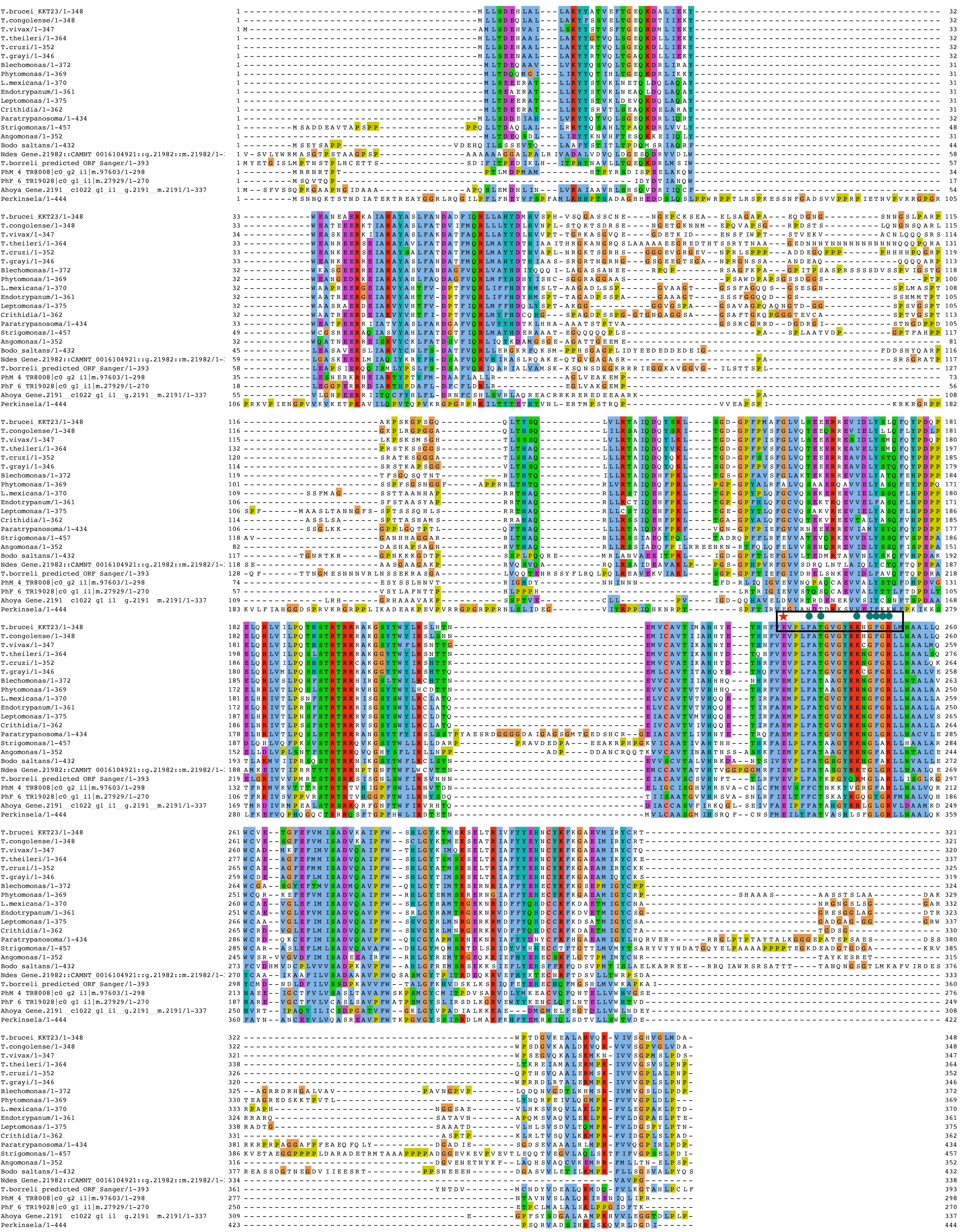
KKT23 is conserved among kinetoplastids. KKT23 has highly conserved protein sequence including the N-terminal helical region and the GNAT domain predicted near the C-terminus (please refer to Figure 1A). The region within 72—126, predicted to be disordered is not conserved (Figure S1B). The active site and residues that form hydrogen bonds with acetyl-CoA are highlighted with red star and teal circles, respectively.

**Figure S4.**
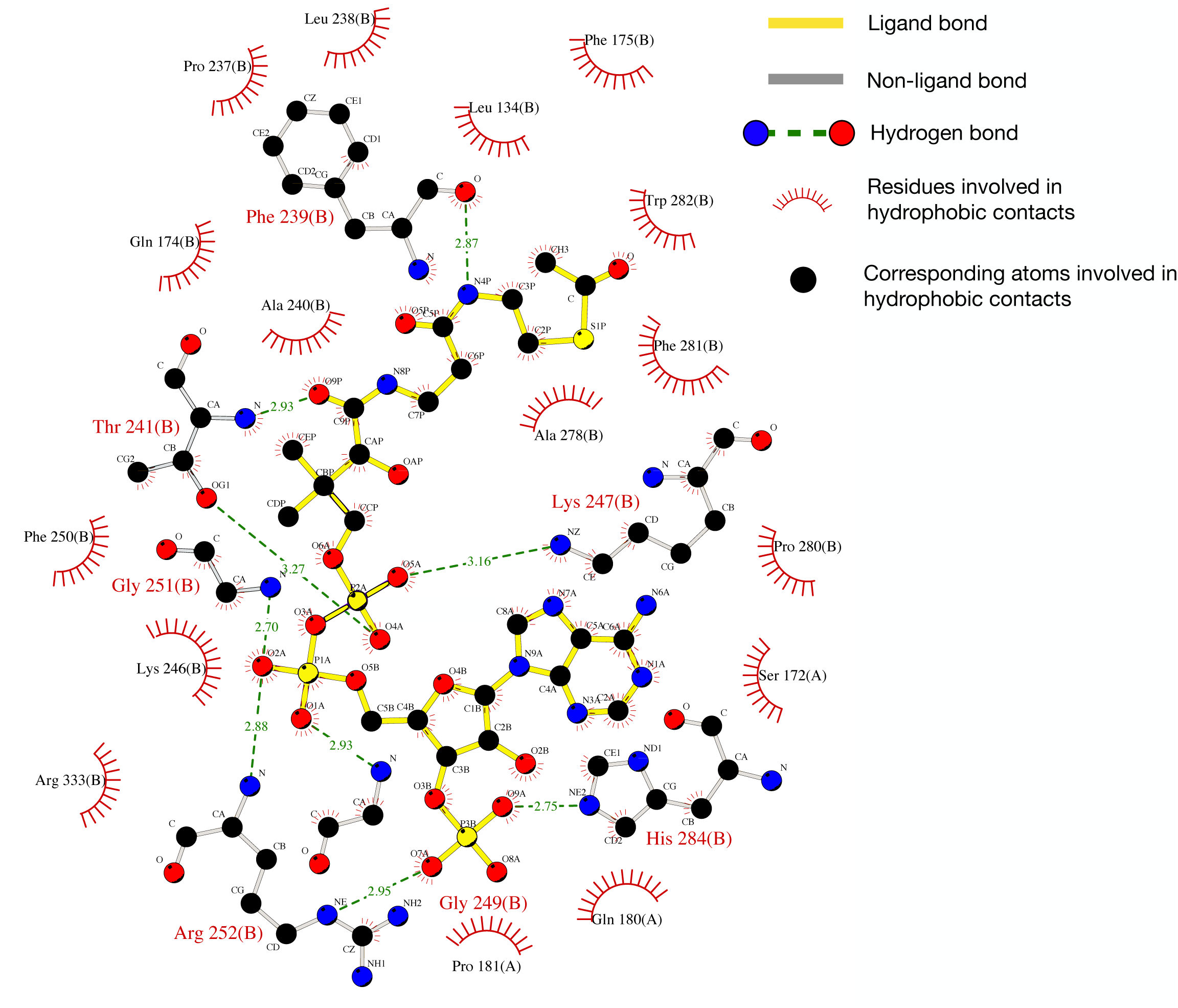
Coordination of acetyl coenzyme A in the crystal structure. This figure shows the visual representation of interactions observed between acetyl-CoA and KKT23 residues in the crystal structure of KKT23^125—348^. The acetyl-CoA is coloured in yellow, while the side chains of KKT23 residues are shown as grey sticks. Acetyl-CoA is coordinated through hydrogen bonds and hydrophobic interactions. The graph shows acetyl-CoA bound to molecule B from the crystal structure but also represents the equivalent ligand in molecule A. This figure was prepared and analysed using LIGPLOT (Wallace et al., 1995).

**Figure S5.**
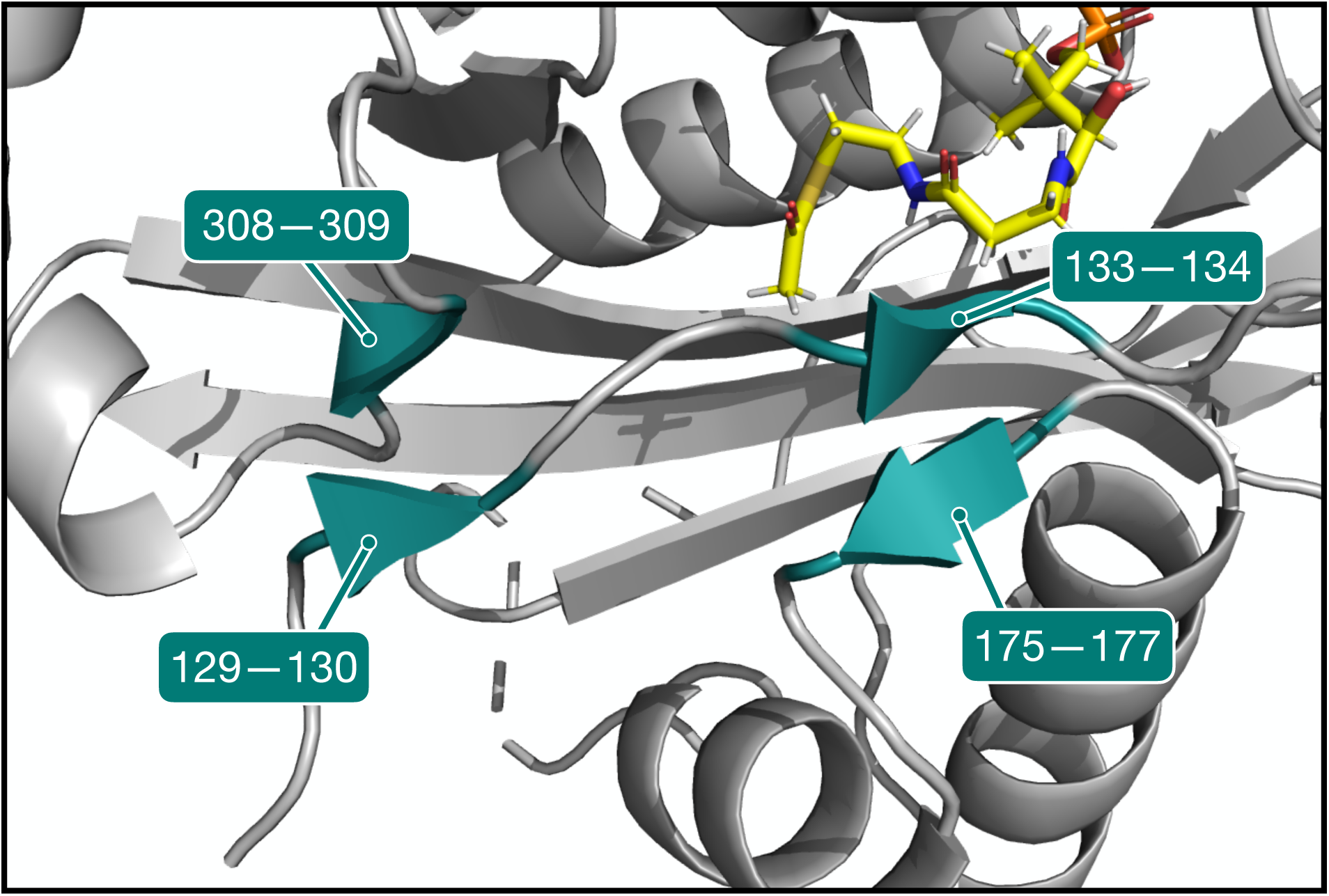
Short β-strands stabilise the unstructured N-terminal region. In the crystal structure of KKT23^125—348^, four short β-strands were identified to stabilise the N-terminal region by forming β-sheets stabilised by hydrogen bonds. The β-strands involving residues 129–130 and 308–309 were not predicted by TALOS-N; this can be explained by missing assignments for residues 130–132 and 307–310. Peaks from these residues may be broadened, indicating conformational exchange of a slower timescale for this β-sheet in solution. TALOS-N does identify region 133–134 to have β-strand propensity but the 175–177 region is not predicted to be β-sheet. Analysis of phi/psi torsion angles in the X-ray structure shows that residues 133–134 have the expected torsion angles for a β-strand while residues 175–177 do not; regular β-strand torsion angles are required to give rise to the chemical shift values that TALOS-N recognises as arising from β-strand structure. The X-ray structure shows a bifurcated hydrogen bond involving 133 CO and both 176 and 177 N, indicating the distorted nature of this short β-sheet. NOESY spectra show NOEs between 177 HN and 133 H⍺ and between 133 HN and 175 H⍺; these NOEs indicate that in solution residues 133–134 and 175–177 are in close proximity as predicted by the X-ray structure.

**Figure S6.**
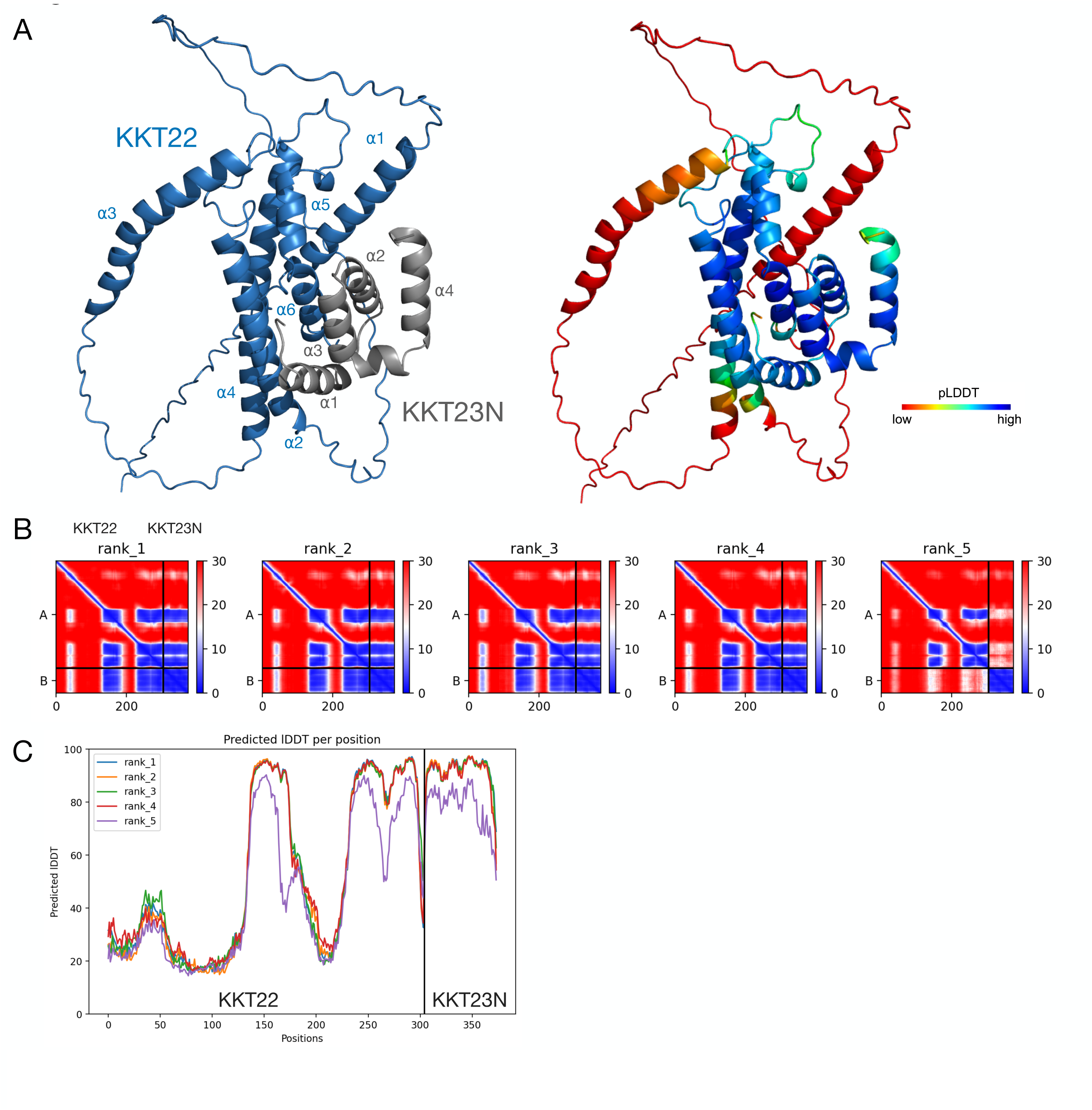
AlphaFold2 model of KKT23N-KKT22 complex. A. AlphaFold2 model (rank 1) of KKT22-KKT23N complex. Cartoon representations are coloured by protein (KKT22 - blue, KKT23N - grey) or according to their pLDDT score. The model was generated over 24 cycles using AF2 colab v1.5.5. The KKT22-KKT23N complex is stabilised by hydrophobic and electrostatic interactions between ⍺2 and ⍺6 on KKT22 and ⍺1—⍺3 on KKT23N. B. Predicted Aligned Error (PAE) plots for five models generated by AlphaFold2, coloured by gradient from red to blue. Red and blue indicate low and high error, respectively. The uncertainty in the predicted distance of two amino acids is colour coded from blue (0 Å) to red (30 Å), as shown on the right bar. C. Predicted Local Distance Test (LDDT) per position for the models (rank 1—5) of KKT22-KKT23N complex, generated by AlphaFold2. Values between 70–90 indicate high accuracy model for the protein backbone.

**Figure S7.**
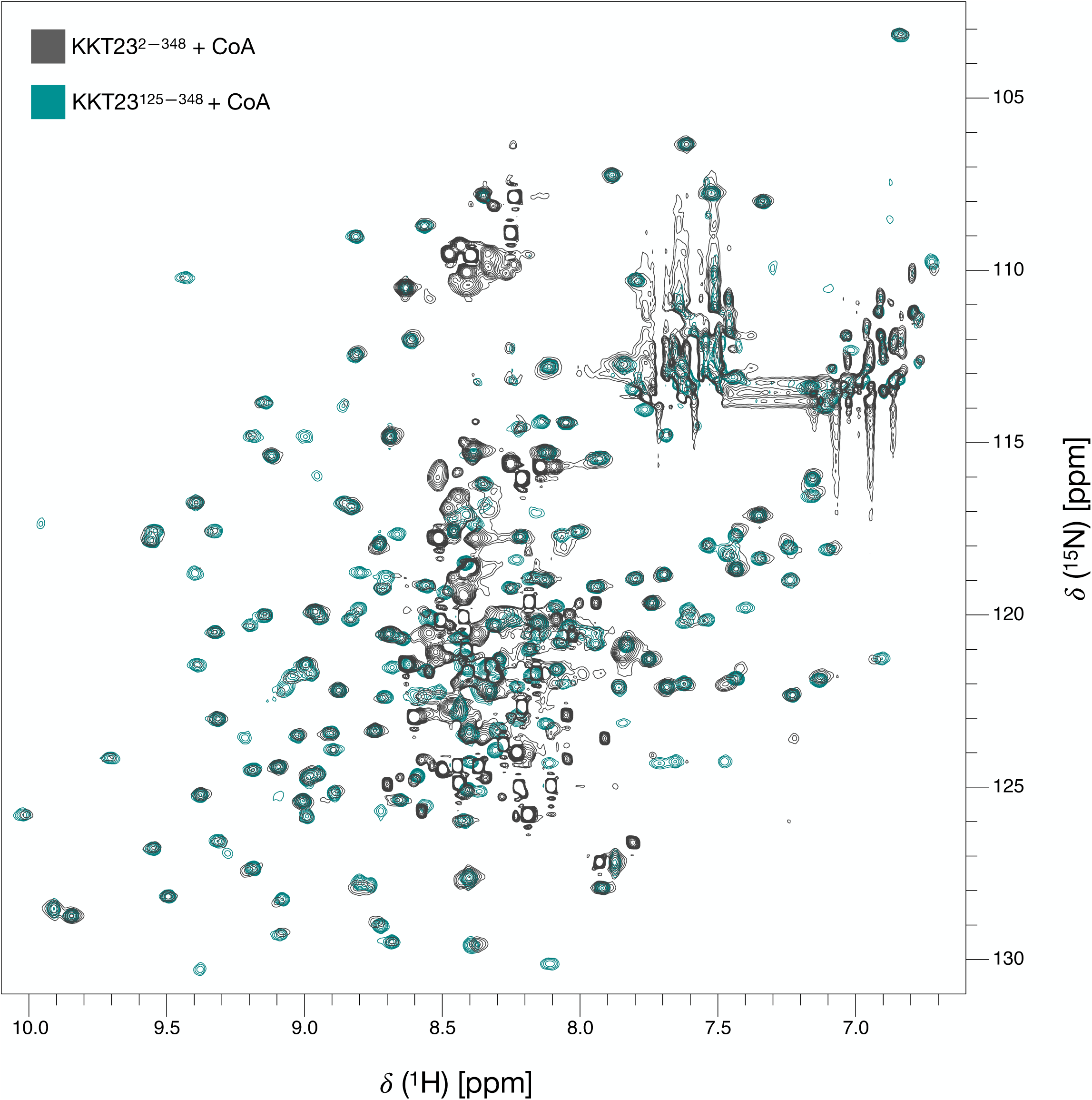
Overlay of spectra obtained for full-length KKT23 and the KKT23 GNAT domain. Overlay of the 750 MHz ^1^H-^15^N BEST TROSY spectra of KKT23^2—348^ (grey) and KKT23^125—348^ (teal). Peaks in the spectrum of KKT23^125—348^ overlay well with the spectrum of KKT23^2—348^, indicating that the properties of the GNAT domain are similar in both constructs. Additional peaks in the spectrum of full-length KKT23 located at ∼8.0–8.7 ppm have high intensity and are not well dispersed, suggesting that they arise from residues that are disordered. Both spectra were collected using KKT23 samples supplemented with coenzyme A.

**Figure S8.**
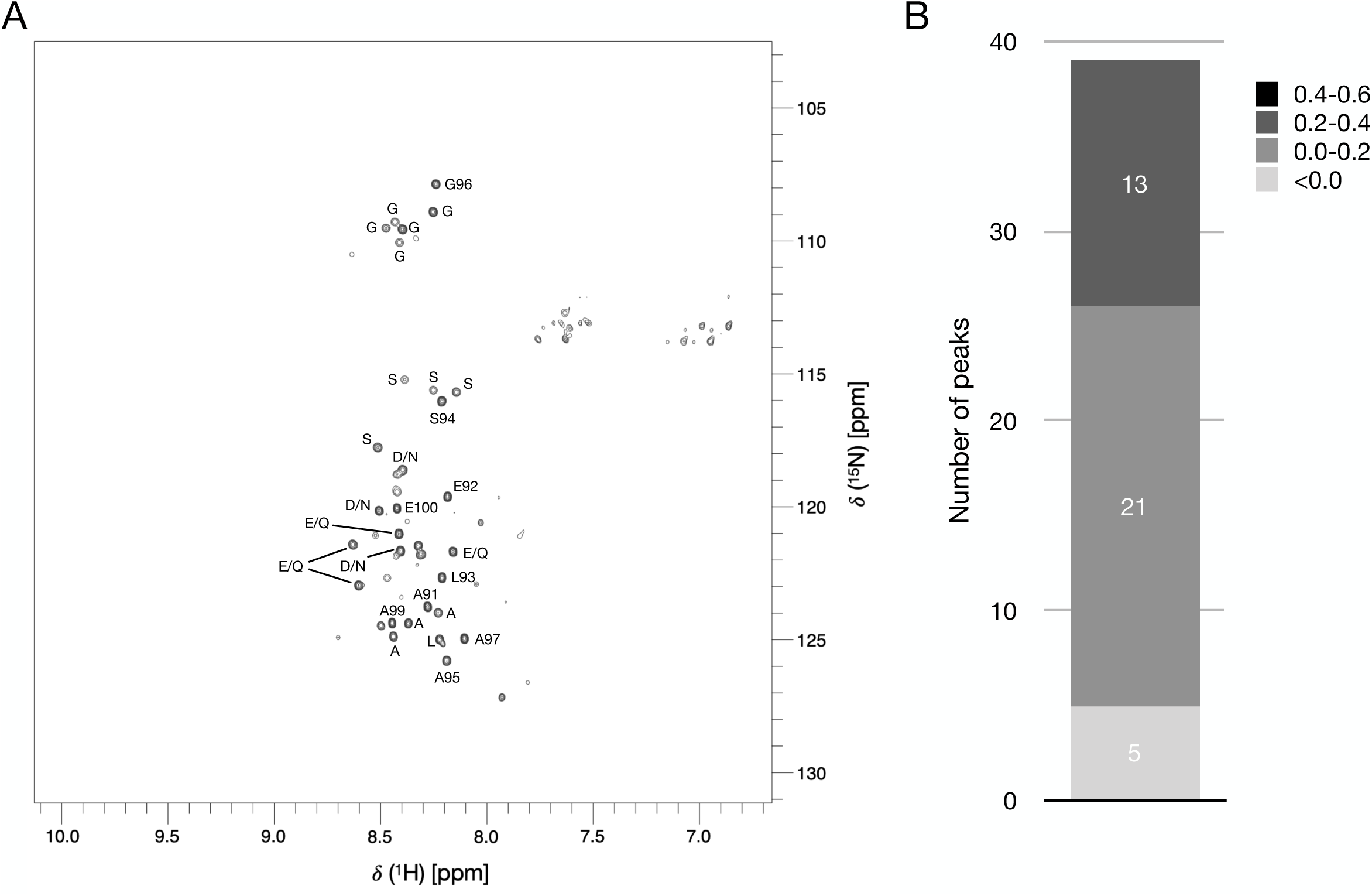
The N-terminal domain and the GNAT domain are connected by a disordered linker. A. The ^1^H-^15^N BEST-TROSY spectrum of the full-length KKT23 in the presence of coenzyme A was contoured in a way to highlight the high intensity peaks around 8.0—8.7 ppm. These peaks likely correspond to disordered regions of KKT23 predicted between 72—126 (Figure S1). Some peaks are labelled with sequence-specific residue assignments or with residue-type information. This information was obtained from analysis of 3D 15N-edited TOCSY-HSQC and NOESY-HSQC spectra. The observed Ala, Gly, Leu, Ser, Asp/Asn and Glu/Gln residues are consistent with the low complexity sequence of the linker region of KKT23. B. The histogram shows the distribution of {^1^H}-^15^N heteronuclear NOE ratios measured in KKT4^2—348^ for 39 of the strongest peaks shown in (A). The heteronuclear NOE ratios were segregated and colour coded by different shades of grey according to different values observed. All peaks displayed the hetNOE ratios <0.4, which is characteristic of a flexible backbone. These peaks likely correspond to residues located within the 72—126 region, which is predicted to be disordered (Figure S1).

**Figure S9.**
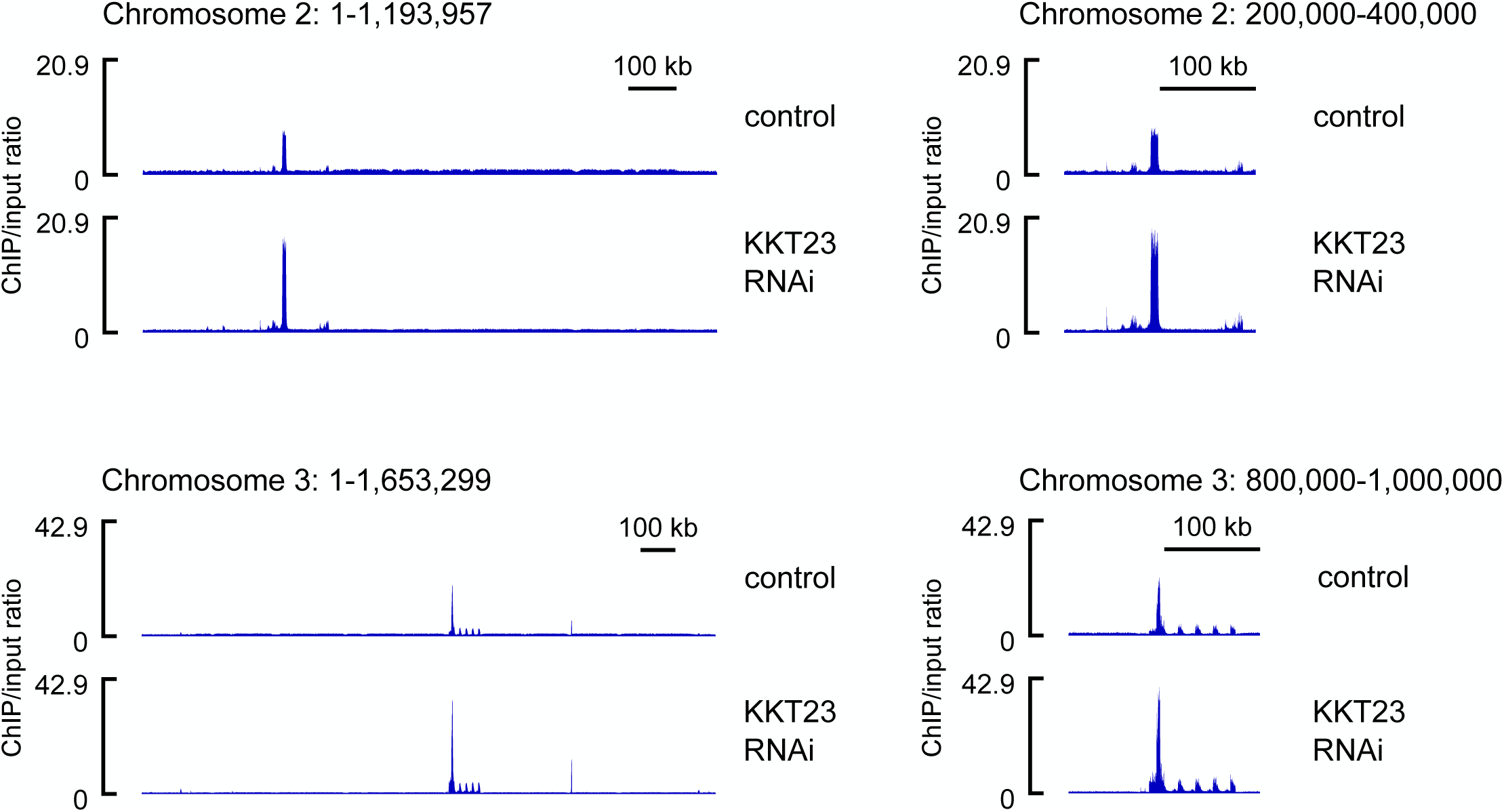
No obvious change in kinetochore positions after KKT23 depletion. ChIP-seq data for KKT3-YFP after depletion of KKT23. RNAi of KKT23 was induced by 1µg/ml doxycycline for 4 days. Data for chromosome 2 (top) and chromosome 3 (bottom) are shown. Cell line, BAP2021

## Key Resources Table

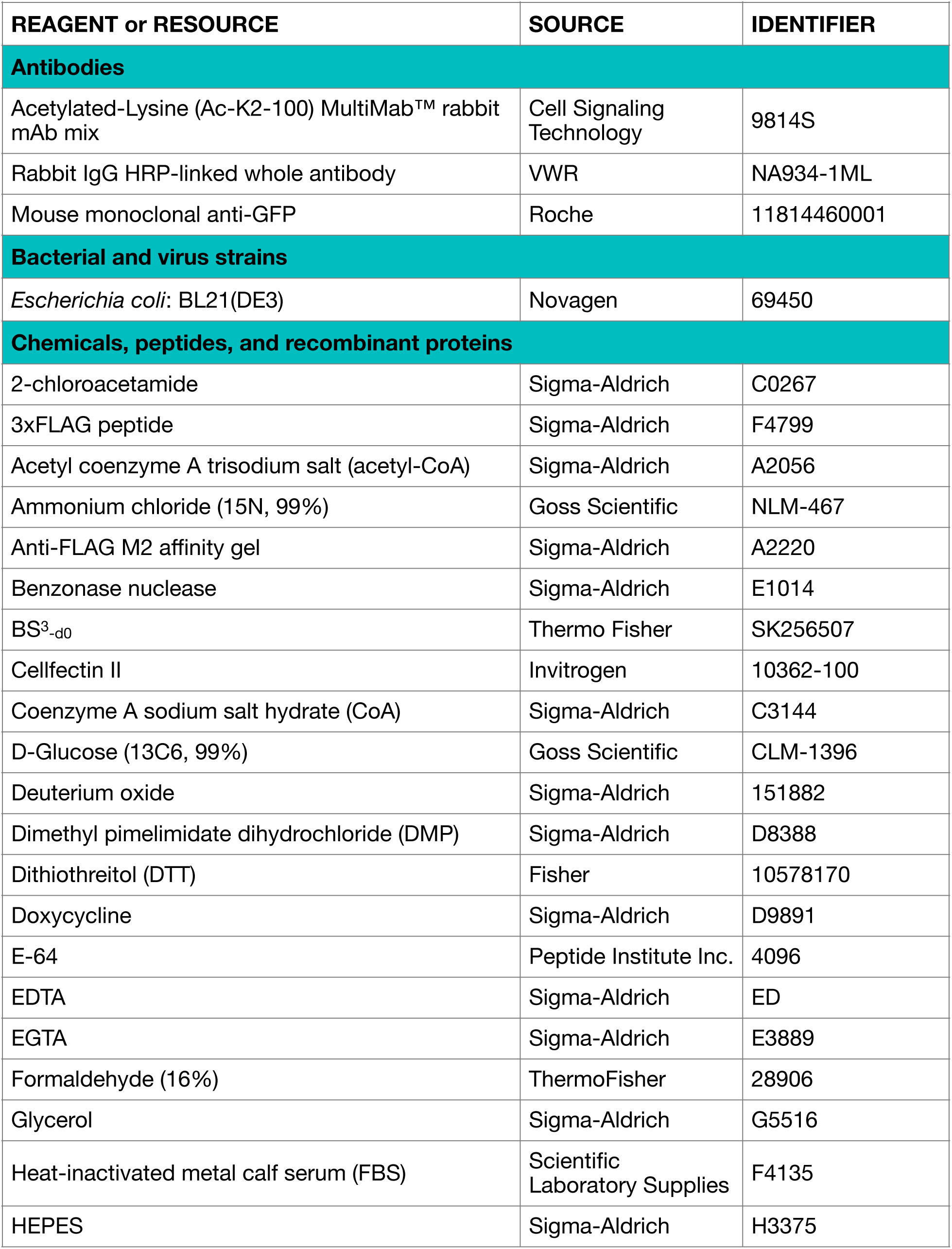
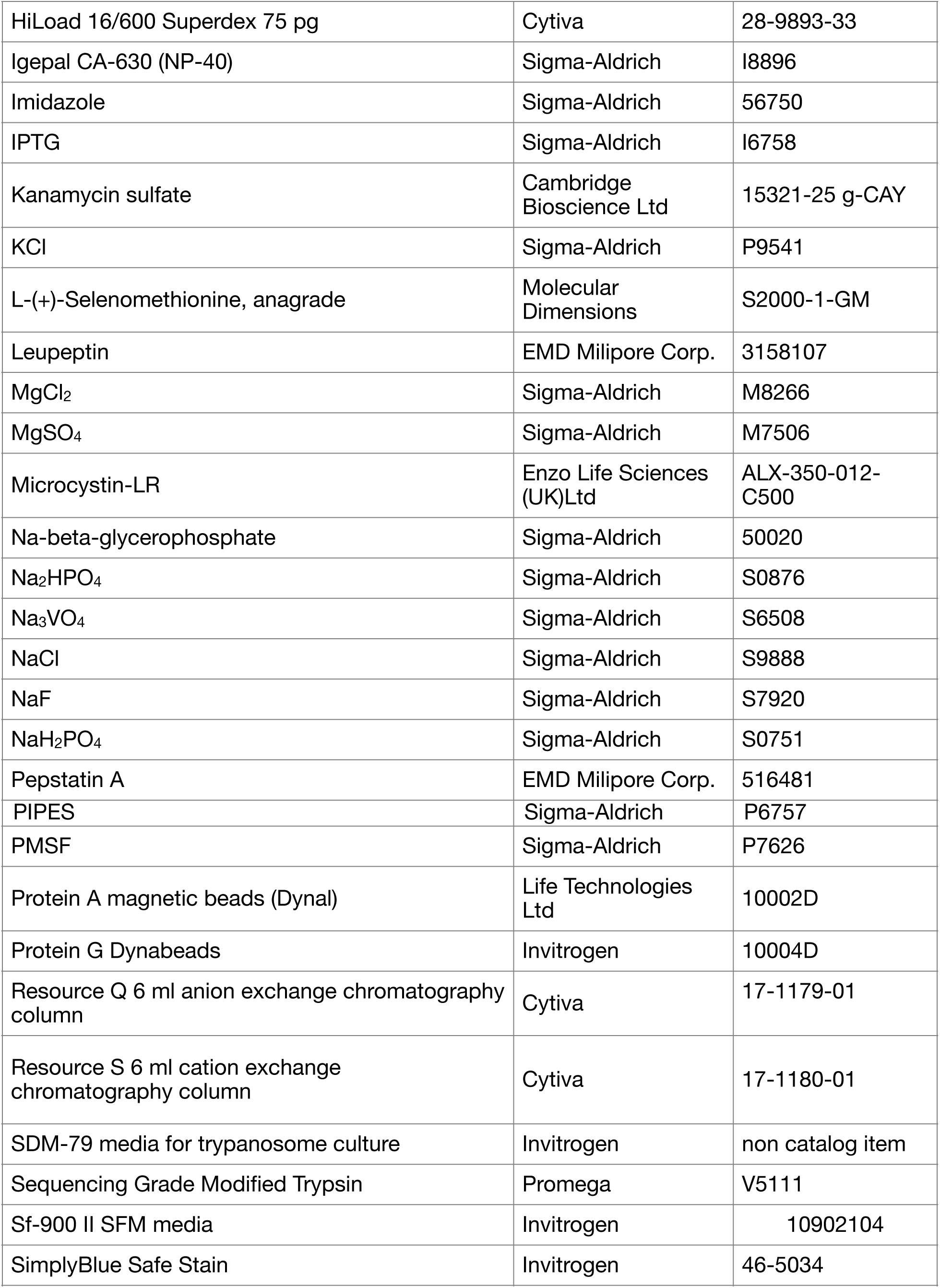
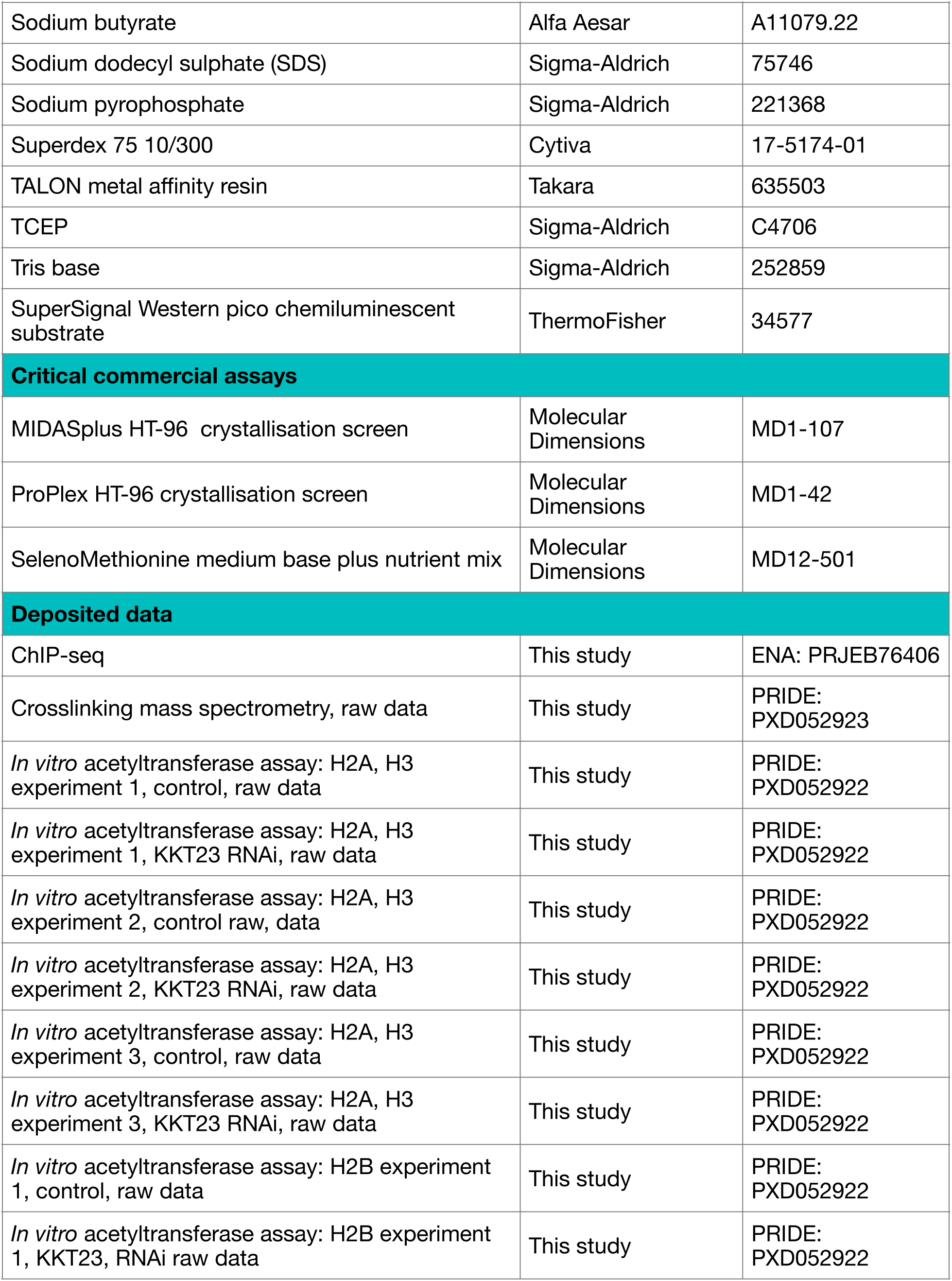
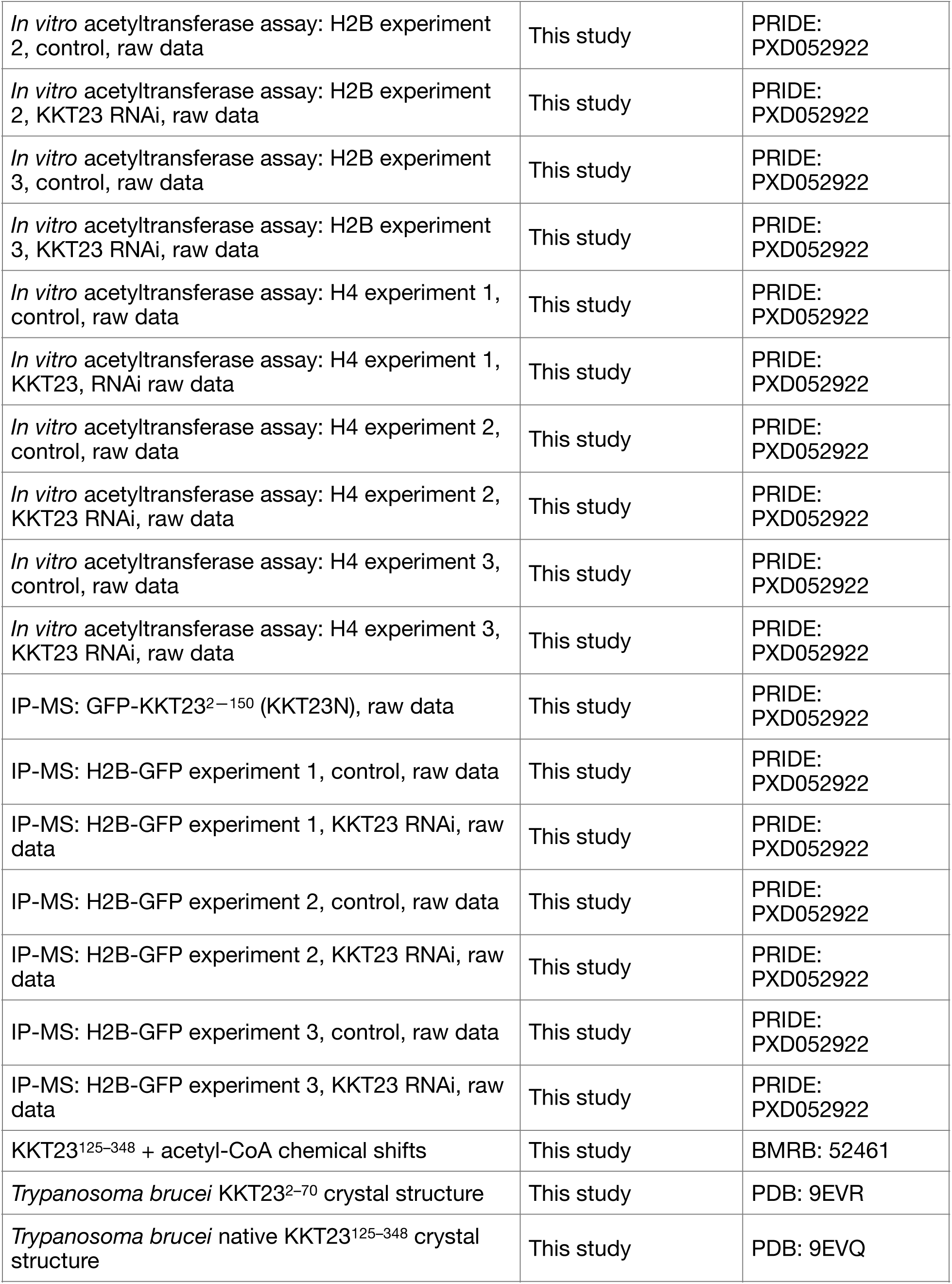
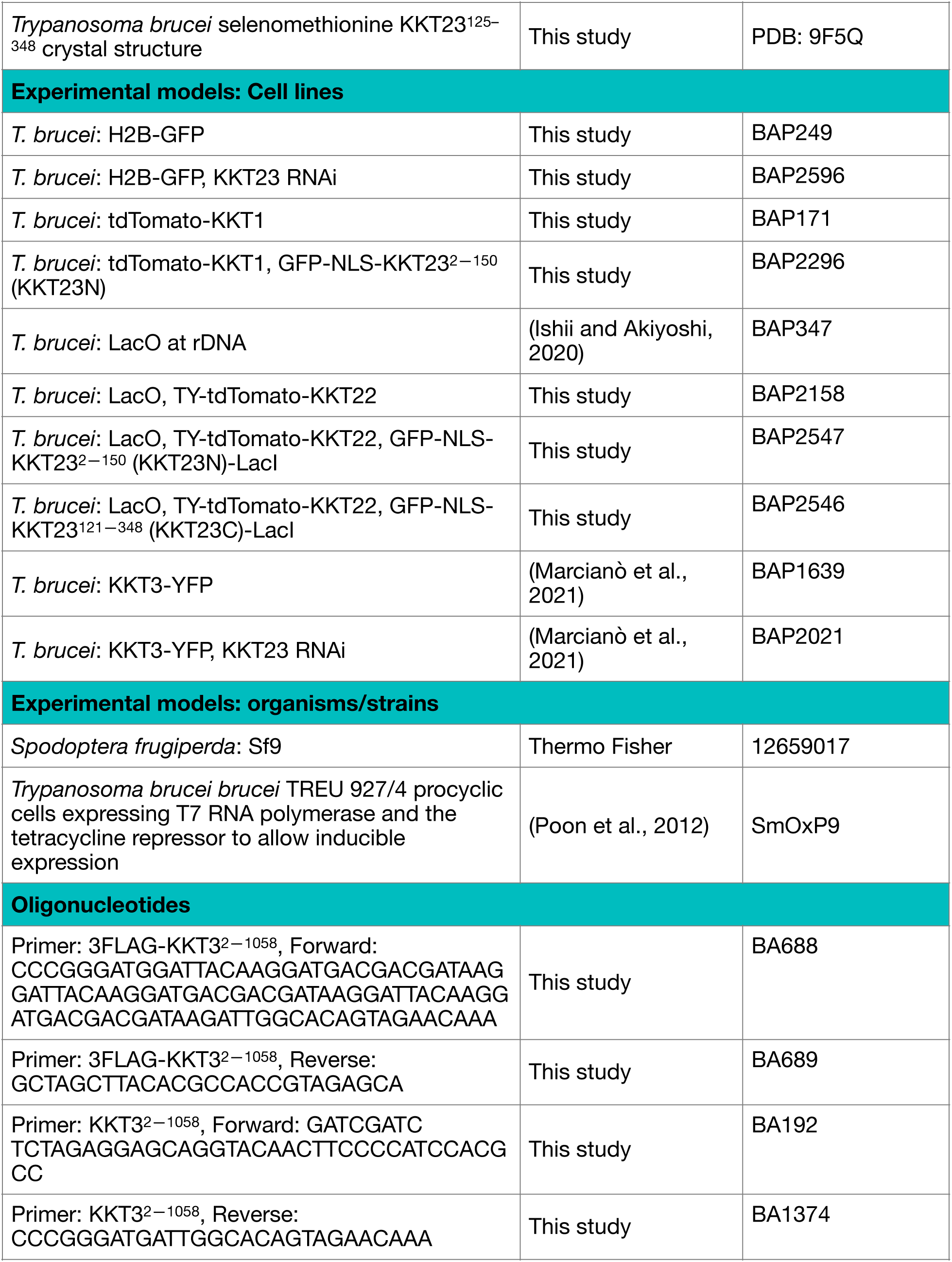
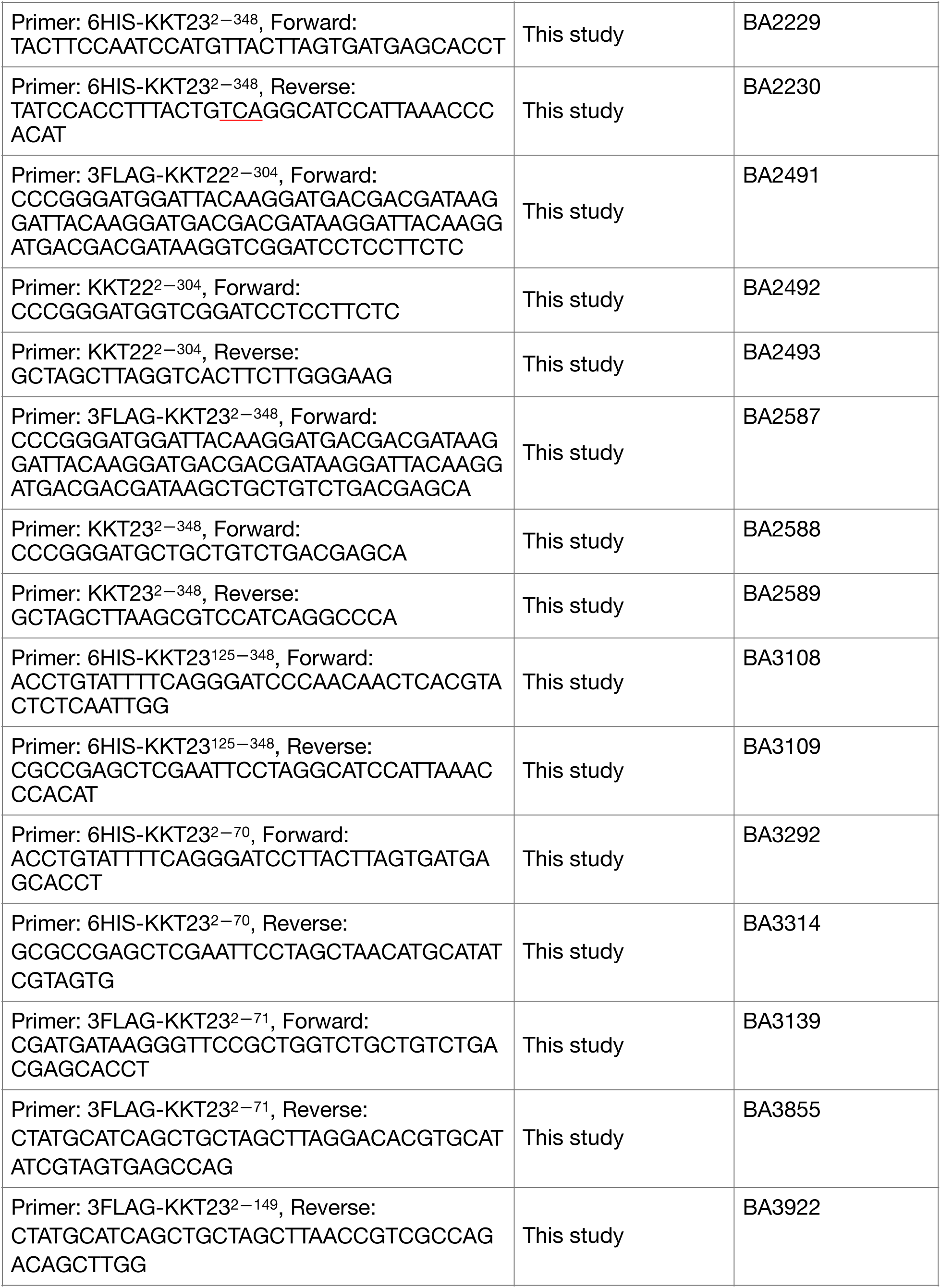
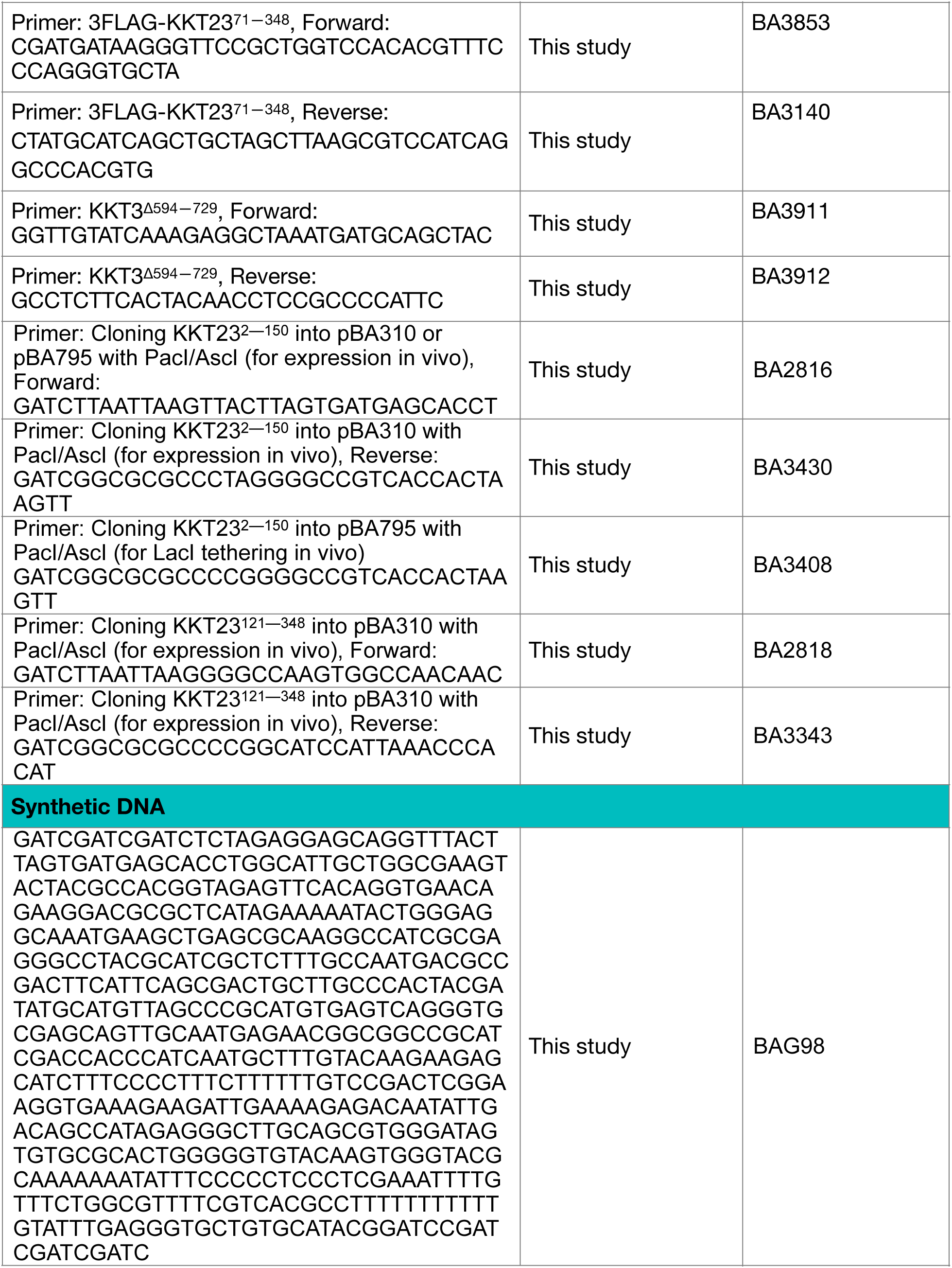
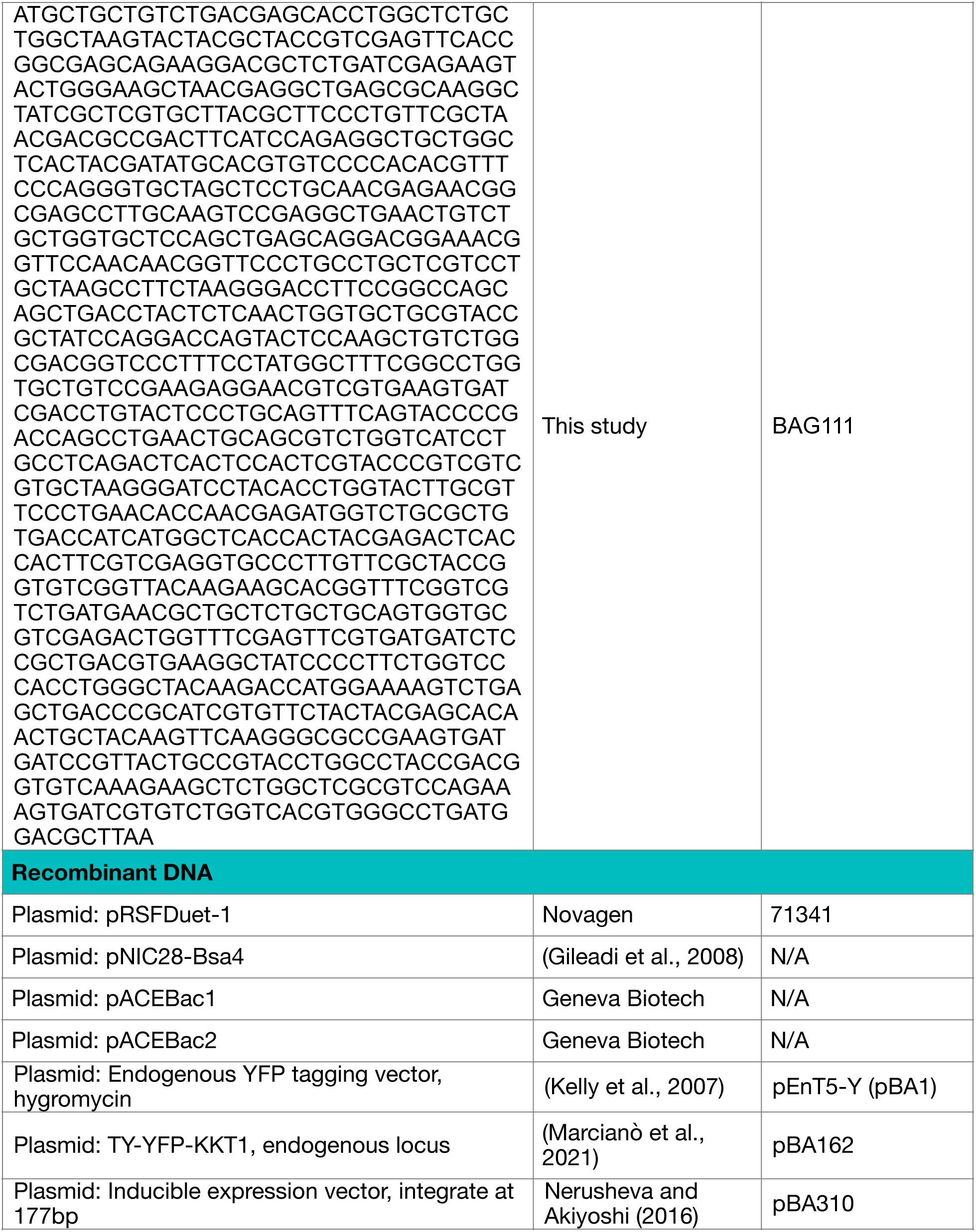
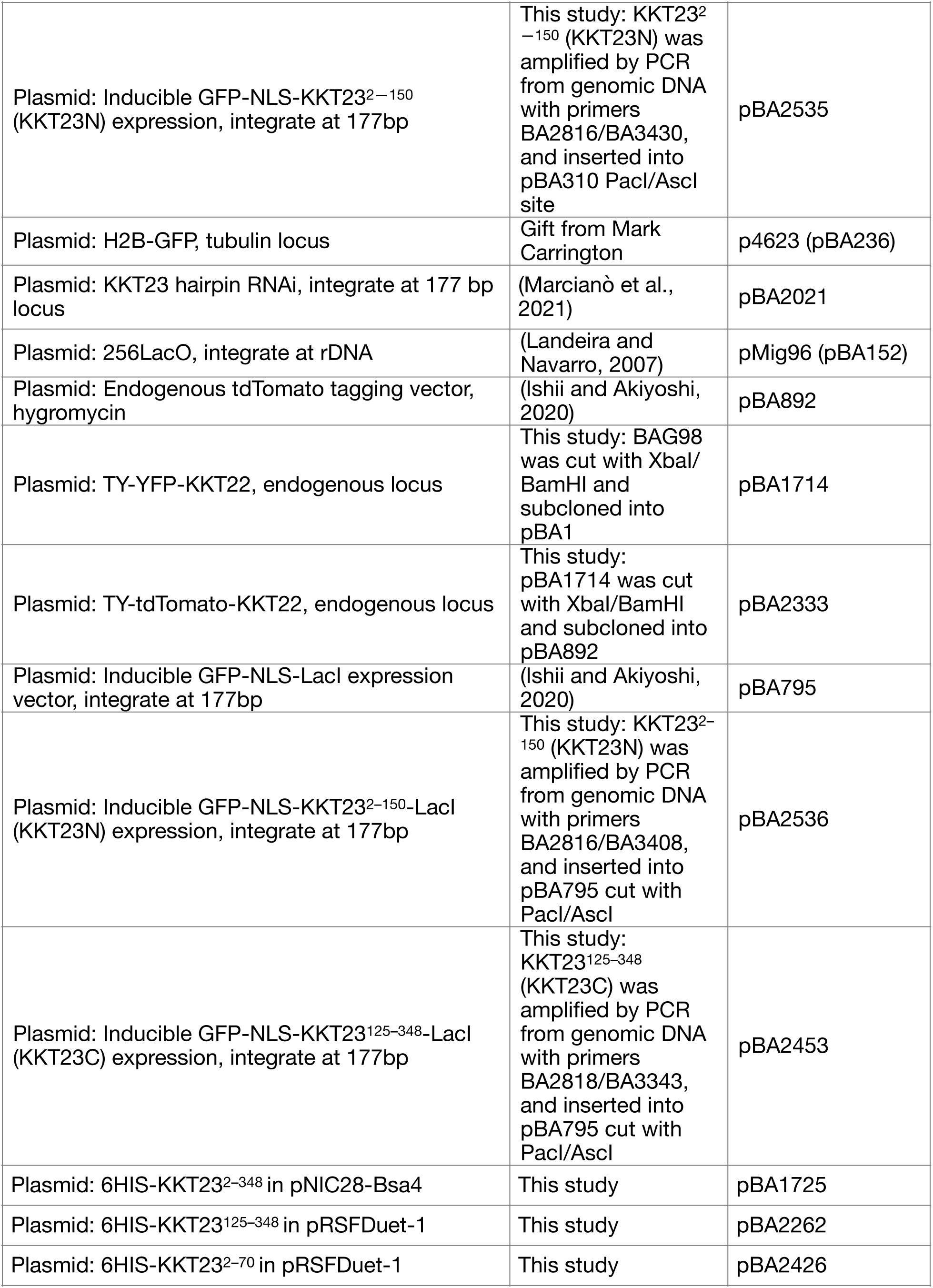
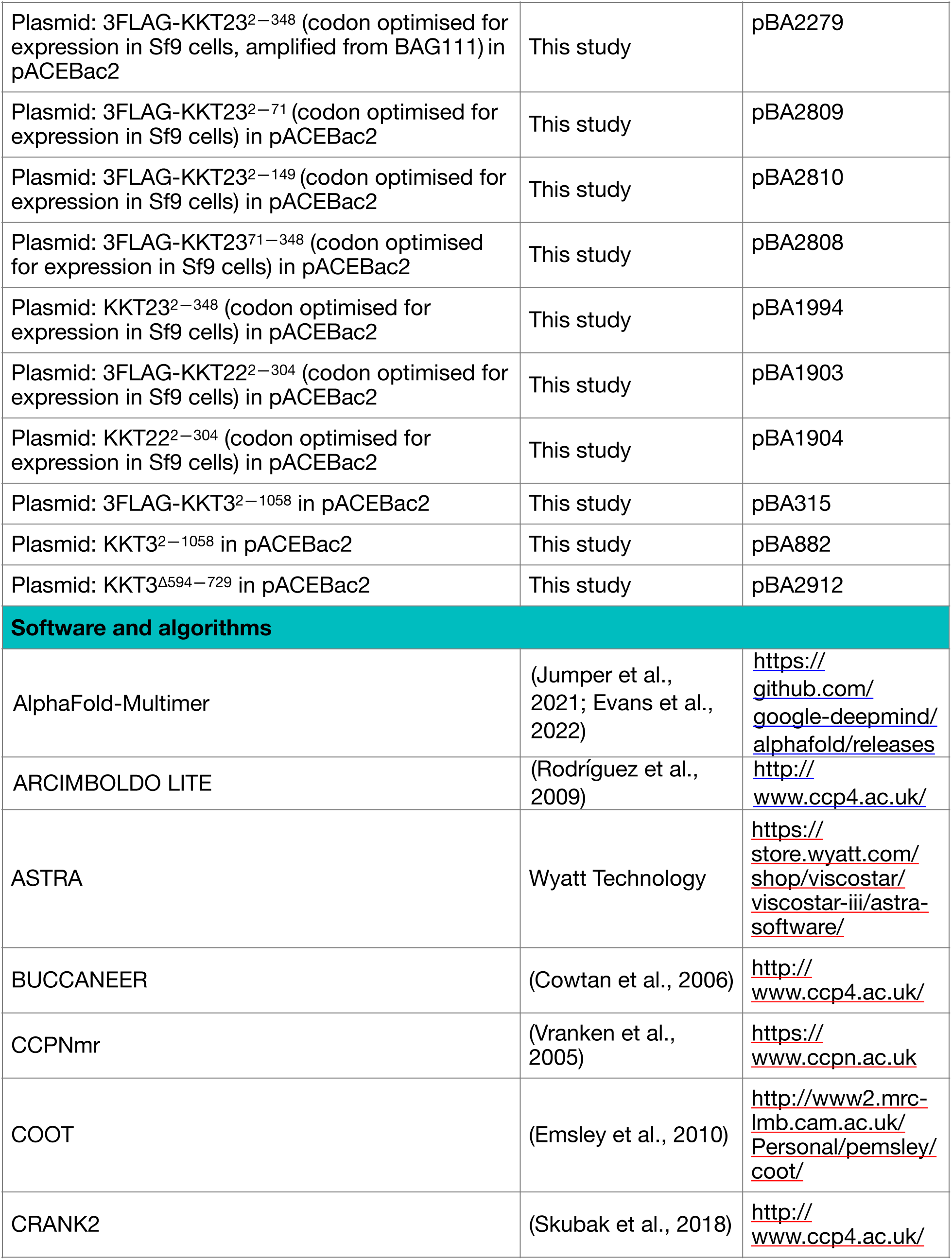
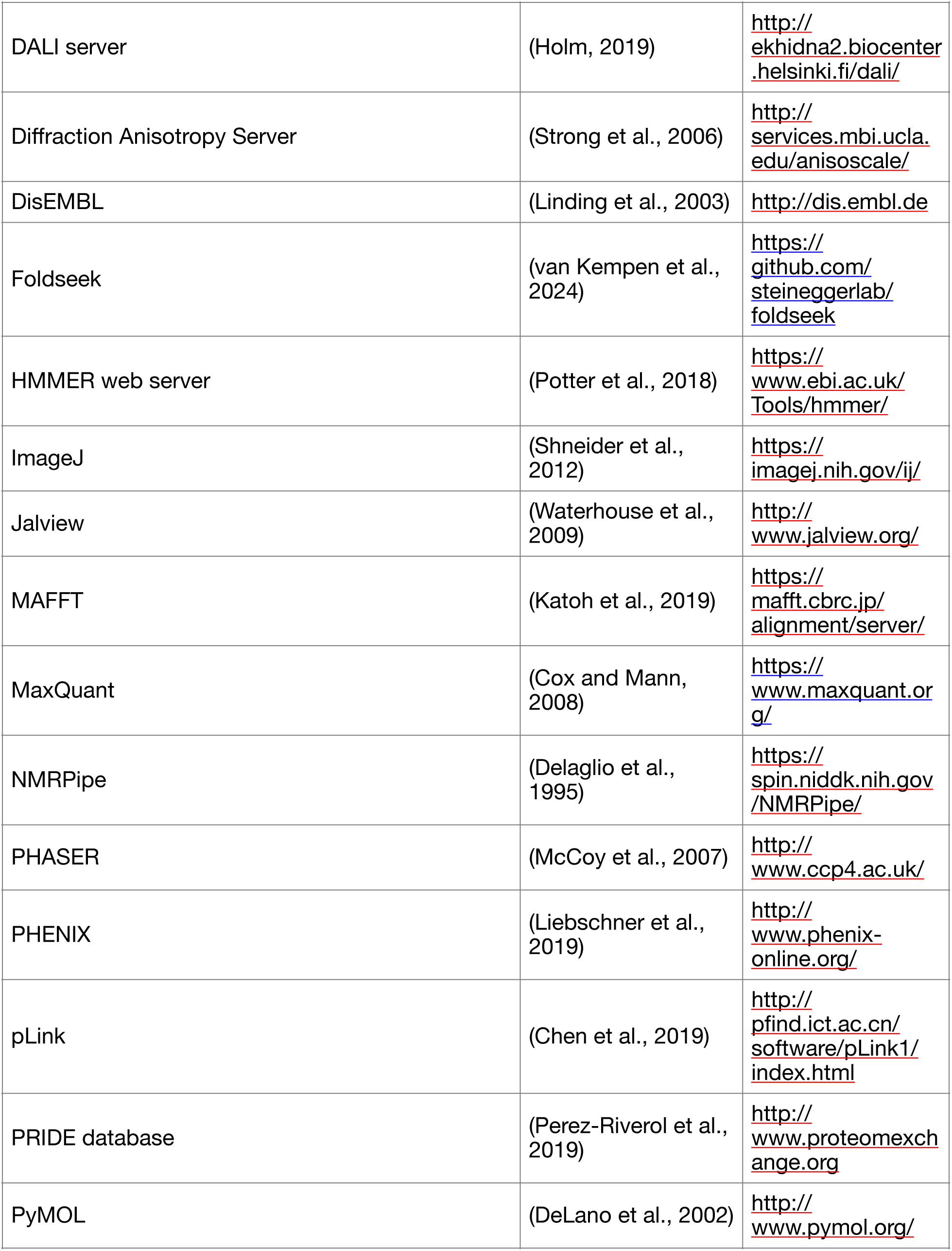
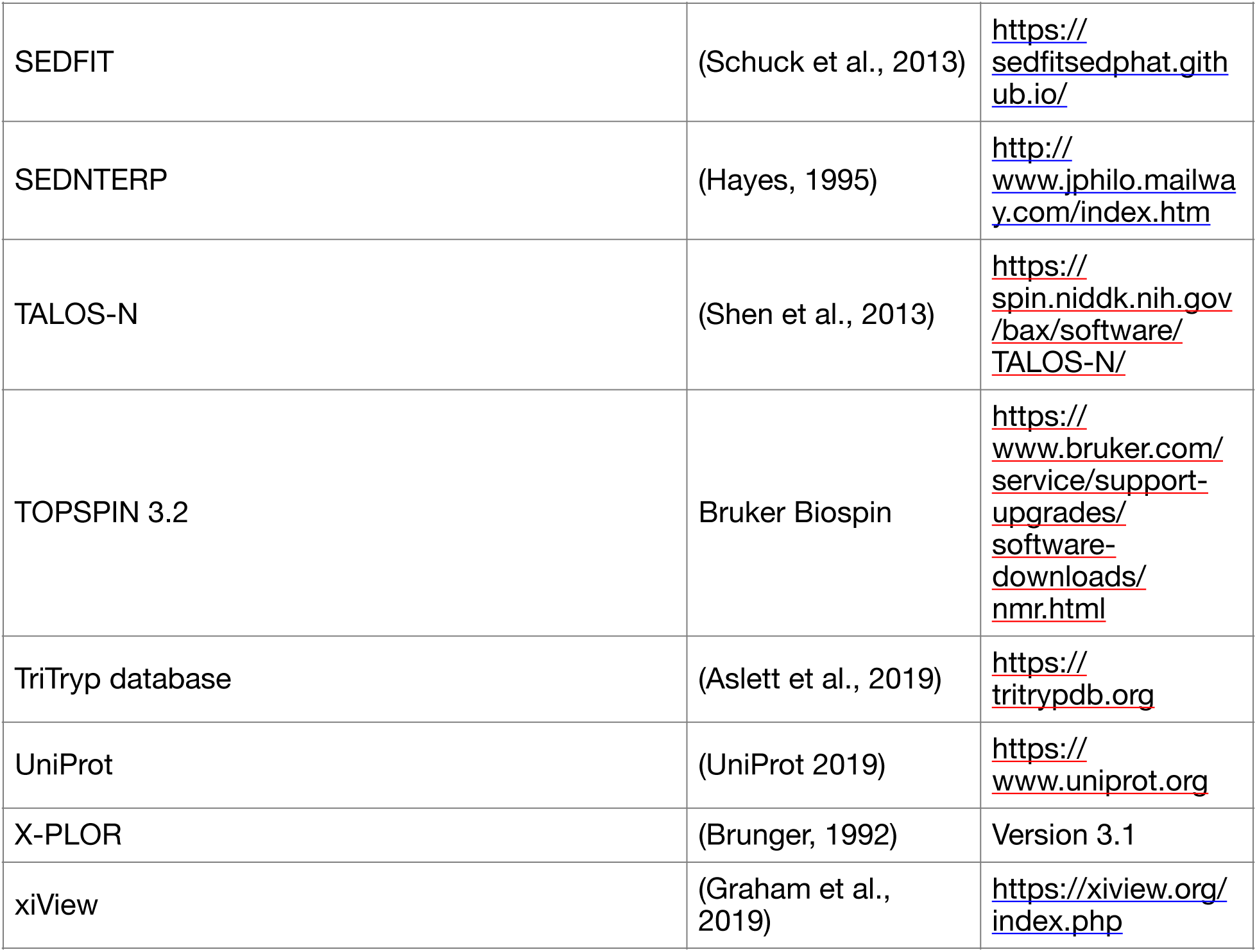

## Bibliography

1. McIntosh, J.R., Mitosis. Cold Spring Harb Perspect Biol, 2016. 8,(9).

2. Cheeseman, I.M., The kinetochore. Cold Spring Harb Perspect Biol, 2014. 6(7): p. a015826.

3. Musacchio, A. and A. Desai, A Molecular View of Kinetochore Assembly and Function. Biology (Basel), 2017. 6,(1).

4. Akiyoshi, B. and K. Gull, Discovery of unconventional kinetochores in kinetoplastids. Cell, 2014. 156(6): p. 1247–1258.

5. Nerusheva, O.O. and B. Akiyoshi, Divergent polo box domains underpin the unique kinetoplastid kinetochore. Open Biol, 2016. 6,(3).

6. Nerusheva, O.O., P. Ludzia, and B. Akiyoshi, Identification of four unconventional kinetoplastid kinetochore proteins KKT22-25 in Trypanosoma brucei. Open Biol, 2019. 9(12): p. 190236.

7. D’Archivio, S. and B. Wickstead, Trypanosome outer kinetochore proteins suggest conservation of chromosome segregation machinery across eukaryotes. Journal of Cell Biology, 2017. 216(2): p. 379–391.

8. Brusini, L., et al., Trypanosome KKIP1 Dynamically Links the Inner Kinetochore to a Kinetoplastid Outer Kinetochore Complex. Front Cell Infect Microbiol, 2021. 11: p. 641174.

9. Marciano, G., et al., Kinetoplastid kinetochore proteins KKT2 and KKT3 have unique centromere localization domains. J Cell Biol, 2021. 220(8).

10. Mo, F., et al., Acetylation of Aurora B by TIP60 ensures accurate chromosomal segregation. Nat Chem Biol, 2016. 12(4): p. 226–32.

11. Ohzeki, J., et al., KAT7/HBO1/MYST2 Regulates CENP-A Chromatin Assembly by Antagonizing Suv39h1-Mediated Centromere Inactivation. Dev Cell, 2016. 37(5): p. 413–27.

12. Dyda, F., D.C. Klein, and A.B. Hickman, GCN5-related N-acetyltransferases: a structural overview. Annu Rev Biophys Biomol Struct, 2000. 29: p. 81–103.

13. DeLano, W.L., Pymol: An open-source molecular graphics tool. CCP4 Newsletter On Protein Crystallography, 2002. 40: p. 82-92.

14. Krissinel, E. and K. Henrick, Inference of macromolecular assemblies from crystalline state. J Mol Biol, 2007. 372(3): p. 774–97.

15. Marmorstein, R. and S.Y. Roth, Histone acetyltransferases: function, structure, and catalysis. Curr Opin Genet Dev, 2001. 11(2): p. 155–61.

16. Shen, Y. and A. Bax, Protein backbone and sidechain torsion angles predicted from NMR chemical shifts using artificial neural networks. Journal of Biomolecular NMR, 2013. 56(**3**): p. 227–241.

17. Kay, L.E., D.A. Torchia, and A. Bax, Backbone Dynamics of Proteins As Studied by 15N Inverse Detected Heteronuclear NMR Spectroscopy: Application to Staphylococcal Nuclease. Biochemistry, 1989. 28(**23**): p. 8972–9.

18. Holm, L., et al., DALI shines a light on remote homologs: One hundred discoveries. Protein Sci, 2023. 32(1): p. e4519.

19. van Kempen, M., et al., Fast and accurate protein structure search with Foldseek. Nat Biotechnol, 2024. 42(2): p. 243–246.

20. Drazic, A., et al., NAA80 is actin’s N-terminal acetyltransferase and regulates cytoskeleton assembly and cell motility. Proc Natl Acad Sci U S A, 2018. 115(17): p. 4399–4404.

21. Goris, M., et al., Structural determinants and cellular environment define processed actin as the sole substrate of the N-terminal acetyltransferase NAA80. Proc Natl Acad Sci U S A, 2018. 115(17): p. 4405–4410.

22. Kuo, M.H., et al., Transcription-linked acetylation by Gcn5p of histones H3 and H4 at specific lysines. Nature, 1996. 383(6597): p. 269–72.

23. Yang, X.J., et al., A p300/CBP-associated factor that competes with the adenoviral oncoprotein E1A. Nature, 1996. 382(6589): p. 319–24.

24. Bu, P., et al., Loss of Gcn5 acetyltransferase activity leads to neural tube closure defects and exencephaly in mouse embryos. Mol Cell Biol, 2007. 27(9): p. 3405–16.

25. Lee, K.K. and J.L. Workman, Histone acetyltransferase complexes: one size doesn’t fit all. Nat Rev Mol Cell Biol, 2007. 8(4): p. 284–95.

26. Deak, G., et al., Histone divergence in trypanosomes results in unique alterations to nucleosome structure. Nucleic Acids Res, 2023. 51(15): p. 7882–7899.

27. Janzen, C.J., et al., Unusual histone modifications in Trypanosoma brucei. FEBS Lett, 2006. 580(9): p. 2306–10.

28. Mandava, V., et al., Histone modifications in Trypanosoma brucei. Mol Biochem Parasitol, 2007. 156(1): p. 41–50.

29. de Jesus, T.C., et al., Chromatin Proteomics Reveals Variable Histone Modifications during the Life Cycle of Trypanosoma cruzi. J Proteome Res, 2016. 15(6): p. 2039–51.

30. Picchi, G.F., et al., Post-translational Modifications of Trypanosoma cruzi Canonical and Variant Histones. J Proteome Res, 2017. 16(3): p. 1167–1179.

31. Kraus, A.J., et al., Distinct roles for H4 and H2A.Z acetylation in RNA transcription in African trypanosomes. Nat Commun, 2020. 11(1): p. 1498.

32. Figueiredo, L.M., G.A. Cross, and C.J. Janzen, Epigenetic regulation in African trypanosomes: a new kid on the block. Nat Rev Microbiol, 2009. 7(7): p. 504–13.

33. Landeira, D. and M. Navarro, Nuclear repositioning of the VSG promoter during developmental silencing in Trypanosoma brucei. J Cell Biol, 2007. 176(2): p. 133–9.

34. Maree, J.P., et al., Trypanosoma brucei histones are heavily modified with combinatorial post-translational modifications and mark Pol II transcription start regions with hyperacetylated H2A. Nucleic Acids Res, 2022. 50(17): p. 9705–9723.

35. Hong, L., et al., Studies of the DNA binding properties of histone H4 amino terminus. Thermal denaturation studies reveal that acetylation markedly reduces the binding constant of the H4 "tail" to DNA. J Biol Chem, 1993. 268(1): p. 305–14.

36. Brower-Toland, B., et al., Specific contributions of histone tails and their acetylation to the mechanical stability of nucleosomes. J Mol Biol, 2005. 346(1): p. 135–46.

37. Bannister, A.J. and T. Kouzarides, Regulation of chromatin by histone modifications. Cell Res, 2011. 21(3): p. 381–95.

38. Talbert, P.B. and S. Henikoff, Transcribing Centromeres: Noncoding RNAs and Kinetochore Assembly. Trends Genet, 2018. 34(8): p. 587–599.

39. Tschudi, C., et al., Small interfering RNA-producing loci in the ancient parasitic eukaryote Trypanosoma brucei. BMC Genomics, 2012. 13: p. 427.

40. Jurrus, E., et al., Improvements to the APBS biomolecular solvation software suite. Protein Sci, 2018. 27(1): p. 112–128.

41. Perez-Riverol, Y., et al., The PRIDE database and related tools and resources in 2019: improving support for quantification data. Nucleic Acids Res, 2019. 47(D1): p. 442–450.

42. Deutsch, E.W., et al., The ProteomeXchange consortium at 10 years: 2023 update. Nucleic Acids Res, 2023. 51(D1): p. D1539–D1548.

43. Brun, R. and Schonenberger, Cultivation and in vitro cloning or procyclic culture forms of Trypanosoma brucei in a semi-defined medium. Short communication. Acta Trop, 1979. 36(3): p. 289–92.

44. Schuck, P., et al., SEDFIT-MSTAR: Molecular weight and molecular weight distribution analysis of polymers by sedimentation equilibrium in the ultracentrifuge. The Analyst, 2013. 139(1): p. 79–92.

45. Hayes D, L.T.P.J., *Program Sednterp: sedimentation interpretation program.* Alliance Protein Laboratories, Thousand Oaks, CA, 1995.

46. Rodríguez, D., et al., Practical structure solution with ARCIMBOLDO. Acta Crystallographica Section D: Biological Crystallography, 2012. 68(4): p. 336–343.

47. Cowtan, K., The Buccaneer software for automated model building. 1. Tracing protein chains. Acta Crystallographica Section D: Biological Crystallography, 2006. 62(9): p. 1002–1011.

48. Cowtan, K., P. Emsley, and K.S. Wilson, From crystal to structure with CCP4, in Acta Crystallographica Section D: Biological Crystallography. 2011. p. 233–234.

49. Adams, P.D., et al., PHENIX: A comprehensive Python-based system for macromolecular structure solution. Acta Crystallographica Section D: Biological Crystallography, 2010. 66(2): p. 213–221.

50. Skubak, P., et al., A new MR-SAD algorithm for the automatic building of protein models from low-resolution X-ray data and a poor starting model. IUCrJ, 2018. 5(Pt 2): p. 166–171.

51. McCoy, A.J., et al., Phaser crystallographic software. Journal of Applied Crystallography, 2007.

52. Liebschner, D., et al., Macromolecular structure determination using X-rays, neutrons and electrons: recent developments in Phenix. Acta Crystallogr D Struct Biol, 2019. 75(Pt 10): p. 861–877.

53. Delaglio, F., et al., NMRPipe: a multidimensional spectral processing system based on UNIX pipes. J Biomol NMR, 1995. 6(3): p. 277–293.

54. Vranken, W.F., et al., The CCPN data model for NMR spectroscopy: development of a software pipeline. Proteins, 2005. 59(4): p. 687–96.

55. Zhu, G., et al., Protein dynamics measurements by TROSY-based NMR experiments. J Magn Reson, 2000. 143(2): p. 423–426.

56. Burdett, H., et al., BRCA1-BARD1 combines multiple chromatin recognition modules to bridge nascent nucleosomes. Nucleic Acids Res, 2023. 51(20): p. 11080–11103.

57. Poon, S.K., et al., A modular and optimized single marker system for generating Trypanosoma brucei cell lines expressing T7 RNA polymerase and the tetracycline repressor. Open Biol, 2012. 2(2): p. 110037.

58. Ishii, M. and B. Akiyoshi, Characterization of unconventional kinetochore kinases KKT10 and KKT19 in Trypanosoma brucei. J Cell Sci, 2020. 133(8).

59. Lopez-Escobar, L., et al., Stage-specific transcription activator ESB1 regulates monoallelic antigen expression in Trypanosoma brucei. Nat Microbiol, 2022. 7(8): p. 1280–1290.

60. Schneider, C.A., W.S. Rasband, and K.W. Eliceiri, NIH Image to ImageJ: 25 years of image analysis. Nat Methods, 2012. 9(7): p. 671–675.

61. Rappsilber, J., M. Mann, and Y. Ishihama, Protocol for micro-purification, enrichment, pre-fractionation and storage of peptides for proteomics using StageTips. Nat Protoc, 2007. 2(8): p. 1896–906.

62. Olsen, J.V., et al., Higher-energy C-trap dissociation for peptide modification analysis. Nat Methods, 2007. 4(9): p. 709–12.

63. Chen, Z.L., et al., A high-speed search engine pLink 2 with systematic evaluation for proteome-scale identification of cross-linked peptides. Nat Commun, 2019. 10(1): p. 3404.

64. Graham, M., et al., xiView: A common platform for the downstream analysis of Crosslinking Mass Spectrometry data. bioRxiv, 2019: p. 561829.

65. Aslett, M., et al., TriTrypDB: a functional genomic resource for the Trypanosomatidae. Nucleic Acids Res, 2010. 38(Database issue): p. 457–462.

66. UniProt, C., UniProt: a worldwide hub of protein knowledge. Nucleic Acids Res, 2019. 47(D1): p. D506–D515.

67. Butenko, A., et al., Evolution of metabolic capabilities and molecular features of diplonemids, kinetoplastids, and euglenids. BMC Biol, 2020. 18(1): p. 23.

68. Eddy, S.R., Profile hidden Markov models. Bioinformatics, 1998. 14(9): p. 755–63.

69. Katoh, K., J. Rozewicki, and K.D. Yamada, MAFFT online service: multiple sequence alignment, interactive sequence choice and visualization. Brief Bioinform, 2019. 20(4): p. 1160–1166.

70. Waterhouse, A.M., et al., Jalview Version 2-A multiple sequence alignment editor and analysis workbench. Bioinformatics, 2009. 25(9): p. 1189–1191.

71. Jumper, J., et al., Highly accurate protein structure prediction with AlphaFold. Nature, 2021. 596(7873): p. 583–589.

72. Evans, R., et al., Protein complex prediction with AlphaFold-Multimer. bioRxiv, 2022: p. 2021.10.04.463034.

73. Mirdita, M., et al., ColabFold: making protein folding accessible to all. Nat Methods, 2022. 19(6): p. 679–682.

